# Stationary distributions of sex allocation at selection-mutation balance

**DOI:** 10.64898/2026.11.13.766474

**Authors:** Matthieu Alfaro, Sylvain Billiard, Lionel Roques

**Author notes:** Authors are listed in alphabetical order.

## Abstract

By predicting evolutionarily stable strategies of resource allocation between female and male functions under different ecological and genetic conditions, sex-allocation theory aims to explain the diversity of reproductive systems within and among species. A fundamental limitation of this theory, however, is that it generally predicts evolutionarily stable discrete distributions of sexual phenotypes. In natural populations, by contrast, sexual phenotypes often exhibit continuous variation, ranging from pure male to pure female individuals. Here, we investigate the form of continuous stationary distributions of sexual phenotypes arising from mutation–selection balance. We extend classical models of sex allocation by incorporating recurrent, infinitesimal mutations, resulting in a nonclassical replicator–mutator partial differential equation. We show that eleven distinct evolutionarily stable states can arise, depending on the mutation regime and the shape of fitness trade-offs between male and female functions. Remarkably, most reproductive systems described in the literature emerge as limiting or stationary distributions of our model. We further show that mutation alone can drive shifts between reproductive systems by recurrently introducing variants with suboptimal resource allocation, thereby preventing populations from remaining indefinitely at classical evolutionarily stable strategies.

## 1 Introduction

Sex ratio and sex allocation theories (Fisher, 1930; Shaw and Mohler, 1953; Charnov et al., 1976; Charnov, 1982) aim to explain the distribution of sexual phenotypes within and among populations and species. Sex allocation theory in particular states that the variation in the production of male or female gametes, and any associated structures involved in mating, fertilization and syngamy, can be explained as a consequence of a set of assumptions: 1) Resources devoted to sexual reproduction are assumed to be fixed, generating a fitness trade-off between male and female functions; 2) Resource allocation is partially heritable and shows genetic variation within populations; 3) Selection favors phenotypes that have the highest lifetime total fitness, i.e. the highest number of offspring produced through male and female functions, which is partly mediated by competition for the access of one type of gametes to the other, thus generating a frequency or density-dependent selection. Under these assumptions, sex-allocation theory can account for major differences among reproductive systems in anisogamous species (Charnov, 1982), including why some plant and animal species are composed of individuals specialized in producing a single type of gamete, whereas in other species individuals change their sex-allocation strategy during their lifetime.

Studies have yielded contrasting and sometimes contradictory results regarding the validity of these three categories of assumptions in both plants and animals. The existence of trade-offs between male and female functions in hermaphrodites is a long-standing debate (reviewed in Schärer 2009; Ashman 2003). However, recent studies describing the genetic architecture of trade-offs in sex allocation (Gerchen et al., 2026) strongly suggest that trade-offs are not merely theoretical constructs but effectively reflect biological constraints rooted in organisms themselves. Regarding the heritability of resource allocations, even though it was generally difficult to conclusively differentiate the relative contributions of genetic and environmental factors in many studies, it has long been clear that traits associated with sex allocation generally have a substantial heritability and are at least partially genetically determined (Delph and Lloyd, 1991; Ashman and Majetic, 2006). In particular, several experimental evolution studies have shown that heritable variation of sex allocation traits and fitness trade-offs was still substantial for trait evolution to occur even after several generations of artificial selection (Mazer et al., 1999; Chen and Pannell, 2025; Gerchen et al., 2026), suggesting a large standing variation in natural populations, or a large number of genes involved where many mutations can occur each generation. Finally, proxies of resource allocation to female or male functions were often shown to be correlated with fitness. For example, it was shown that the number of seeds produced and seeds sired were positively associated to the number of flower per plant in the hermaphroditic herb *Asclepias exaltata* (Broyles and Wyatt, 1990), or to the number of pistil or stamens in *Pulsatilla alpina* (Chen and Pannell, 2025). Similar results were obtained in plant species with separate sexes (dioecy), for example in *Silene dioica* in experimental (Barbot et al., 2022) or natural conditions (Jolivel et al., 2026).

Even though there is now a relative consensus about the validity of the assumptions underlying sex allocation theory, several observations are not explained. In particular, many species show a large heritable variation in allocation to male and/or female functions, suggesting a continuous rather than a discrete distribution of sexual phenotypes. Delph and Lloyd (1991) for example showed in the gynodi-oecious shrub *Veronica subalpina*, where hermaphrodites and females coexist, that the mean fruit-set of hermaphrodites was significantly lower than that of females. At the same time, the fruit-set production among hermaphrodites varied by several orders of magnitude. This resulted in overlapping distributions of allocation to female functions between hermaphrodites and pure females (Fig. 1a). Delph and Lloyd (1991) further showed that this large variation among hermaphrodites was not due to pollen limitation, that it was consistent within individuals across years, and that closely related genotypes tend to have similar resource allocation, strongly suggesting a genetic basis underlying the variation in sex allocation. Similarly, Saumitou-Laprade et al. (2018) estimated the distribution of sex allocation in the common ash tree *Fraxinus excelsior* as the proportion of male, female or hermaphroditic flowers produced in a year by different individuals. They showed a quasi-continuous distribution ranging from pure males to almost pure females within two populations (Fig. 1b). They also showed that the sex allocation was consistent within individuals and between years, and among clones. In another example, Gerchen et al. (2026) artificially removed males in experimental populations of *Mercurialis annua* taking the opportunity of heritable leakiness of females. They showed that the populations rapidly evolved such that females differentially allocated resources in male functions, generating after a few generations a continuous distribution of relative male reproductive effort (Fig. 1c). Finally, Maurice and Fleming (1995) used seed and pollen production to define sexual function of individuals from several populations. They found a bimodal distribution spanning from pure males to pure females (Fig. 1d). Many other studies also showed a large variation in male and/or female allocation among hermaphroditic individuals across years (e.g. Mazer et al. 1999; Laugier et al. 2023; Chen and Pannell 2025). Even in dioecious species, it has been shown that pure males or pure females can be inconstant and produce organs of the opposite sex (Ehlers and Bataillon, 2007; Cossard and Pannell, 2019).

**Figure 1:**
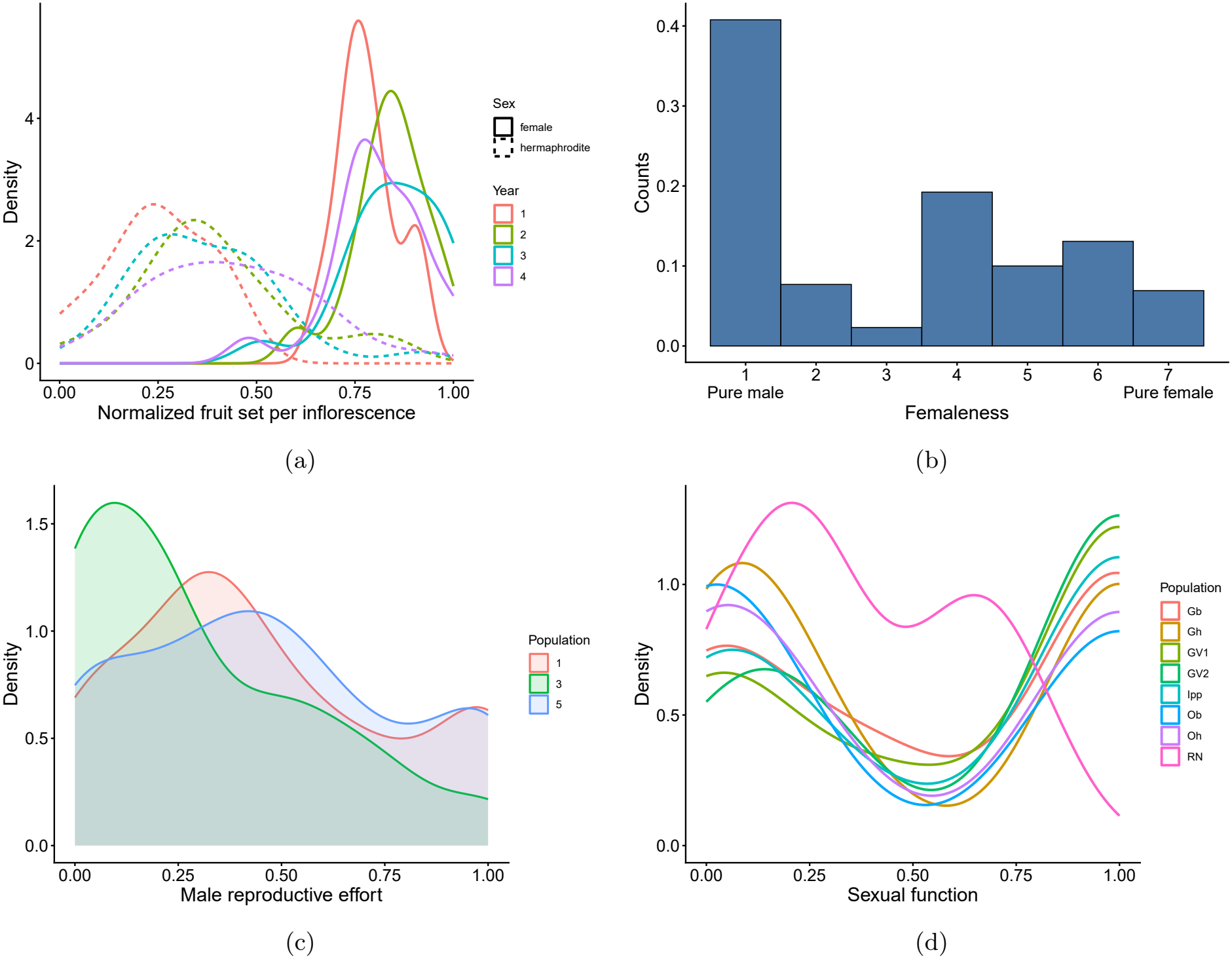
Examples of empirical distribution of sex allocation. (a) Distribution of fruit set per inflorescence on hermaphroditic or female individuals of the perennial plant species *Veronica subalpina* on four successive years (data collected from Fig. 1 in Delph and Lloyd 1991); (b) Distribution of the seven classes of individuals in *Fraxinus excelsior* defined in Saumitou-Laprade et al. (2018) depending on the proportion of male vs. female flowers produced by an individual within a year; (c) Distribution of the relative male reproductive effort from experimental populations of *Mercurialis annua* (data from Gerchen et al. 2026); (d) Distribution of sexual function (*sensu* Lloyd 1980) in *Silene acaulis* (data from Maurice et al. 1998).

The existence of such substantial heritable variation in sex allocation along a continuum from pure females to pure males raises the question of the form of the stationary distribution expected at mutation-selection equilibrium. This question is even more intriguing because there is a striking discrepancy with predictions from the current sex allocation theory as it predicts the existence of only one or two coexisting phenotypes within a population. For example, Charnov et al. (1976) and Lesaffre et al. (2024) derived evolutionarily stable strategies (ESSs) in which either a single hermaphroditic phenotype, a hermaphroditic phenotype coexisting with pure males or females, or both pure males and pure females coexist. These predicted ESSs cannot explain the broad, often continuous, distributions of sex allocation observed within populations, nor the substantial variation among individuals that can persist over time and show evidence of heritability. In particular, they do not account for the possibility that multiple phenotypes spanning a continuum of sex allocation strategies may coexist at evolutionary equilibrium. This discrepancy raises the broader question of how selection and mutation shape the stationary distribution of sex allocation, and, in particular, under what conditions different forms of stationary distributions can emerge. The answer to this question is not trivial as mutation, by regularly introducing new sexual phenotypes, will affect selection on sex allocation, as selection on allocation is frequency dependent.

Continuous-trait mutation-selection models based on partial differential equations (PDEs) or integro-differential equations provide a classical framework for describing the evolution of phenotypically structured populations under the joint effects of selection and recurrent mutation (Kimura, 1965; Bürger, 1986; Champagnat et al., 2006, 2008; Alfaro and Carles, 2014, 2017; Martin and Roques, 2016; Gil et al., 2019). In these models, mutations are represented by diffusion or more general operators in an abstract phenotypic space, whereas selection acts through phenotype-dependent fitness or growth terms. This framework, developed mostly for asexual populations, has been used to investigate the dynamics of fitness distributions, adaptation towards phenotypic optima, evolutionary branching, and adaptation in multidimensional or heterogeneous fitness landscapes (Alfaro and Veruete, 2019; Hamel et al., 2020, 2021; Mirrahimi and Gandon, 2020; Alfaro et al., 2023, 2025). Sexual reproduction with continuous traits and recurrent mutation can alternatively be described within quantitative-genetics frameworks, notably through the infinitesimal model (Barton et al., 2017), as well as through related mathematical models explicitly accounting for Mendelian diploid inheritance or nonlocal sexual reproduction (Collet et al., 2013; Dekens and Mirrahimi, 2022). These approaches provide a more explicit description of inheritance and sexual reproduction. However, their mathematical structure differs substantially from standard replicator-mutator equations because reproduction itself couples parental trait distributions through a nonlocal operator. Here, we adopt a complementary approach by formulating the evolutionary dynamics at the level of transmitted gene copies rather than diploid individuals. This allows us to retain the tractable structure of a replicator-mutator PDE while accounting for sexual reproduction through a frequency-dependent fitness functional. In contrast with classical asexual mutation-selection models, fitness is not prescribed independently of the population distribution: because male gametes compete for the fertilization of female gametes, the fitness of a trait emerges from population-wide quantities determined by the current distribution of sex-allocation strategies.

Our goal in this paper is to study the stationary distributions of the resources allocated to male or female gametes production in a population of an anisogamous species at mutation-selection balance. Our model extends classical sex-allocation models (Charnov et al., 1976) and their more recent developments (Lesaffre et al., 2024) by explicitly incorporating recurrent mutation by means of a structured partial differential equation modeling approach. It encompasses continuous stationary distributions under recurrent mutation and, in the limit where mutation becomes weak relative to selection, discrete distributions that can be related to classical ESS predictions. We first show the different possible stationary distributions that can be obtained asymptotically across weak to strong mutational diffusion regimes for different forms of the trade-off functions. Second we use numerical approaches to determine how the regimes of mutation and the shape of the trade-offs affect stationary distributions. Globally, we show that the possible qualitatively different stationary distributions are numerous. Some of the predicted continuous stationary distributions are in addition compatible with observations and thus resolve the apparent discrepancy between the predictions by sex allocation theory and observations.

## 2 Model

### 2.1 Assumptions

We denote by *x* ∈ [0, 1] the proportion of resource allocated to the female function by an individual. Our main assumptions are the following:

1. **Rates of gamete production**. Individuals investing a proportion *x* of resources in female function yield *K*_*f*_ (*x*) female gametes per unit of time and *K*_*m*_(*x*) male gametes per unit of time. The function *K*_*f*_ is increasing and the function *K*_*m*_ is decreasing, which reflects that individuals suffer from a reproductive trade-off because of the fixed amount of resources they can allocate to either male or female gametes. The functional forms of *K*_*f*_ and *K*_*m*_ will be specified later when needed.
2. **Male gametes are not limiting**. All ovules produced in the population are fertilised. Ovules are the only gametes limiting the reproduction rate (no male gamete limitation).
3. **Mating is panmictic**. The rate of mating between individuals does not depend on their trait *x*. Let 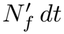 and 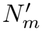 *dt* be, respectively, the numbers of ovules and male gametes released in the population during a very short time *dt* ≪ 1. A male gamete is thus successful in fertilizing an ovule with probability 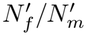. Note that Assumption 2 ensures that 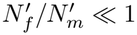.
4. **Selection and fitness**. Fitness is measured as the rate of *haploid* gene copies transmitted to zygotes by an individual with trait *x*. Each ovule transmits a single gene copy from the maternal parent, while each successful male gamete transmits a single gene copy from the paternal parent.
5. **Mutation**. Mutations are assumed to be recurrent, with small and unbiased effects on the trait *x* in the interior of the trait space. Their cumulative effect is modeled by diffusion with coefficient *D >* 0. As *x* ∈ [0, 1], we impose reflecting boundary conditions, which prevent mutational flux from leaving the biologically admissible trait space.

### 2.2 Derivation of a replicator-mutator PDE

Let *u*(*t, x*) denote the *distribution of gene copies* (or “allelic distribution”) at time *t* that encodes the trait value *x* controlling resource allocation to either male or female gametes production. It is *not* the distribution of individual organisms with trait *x*. The paternal and maternal contributions to a zygote are counted separately, each in the relevant *u*(*t, x*) class.

We define the total fitness *K*[*u*](*t, x*) as the rate at which haploid sets are transmitted. The notation [*u*] highlights that fitness of a particular given trait *x* depends on the whole distribution *u*. Under Assumptions 1–4, the expected genetic contribution of an *x*-individual is

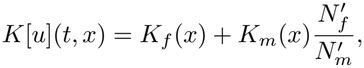

*i*.*e*. the sum of its contribution through the female and male functions. For the sake of simplicity, the notation [*u*] is dropped in the following and we write *K*(*t, x*) instead of *K*[*u*](*t, x*), the dependence on *u* remaining implicit.

With the notation above

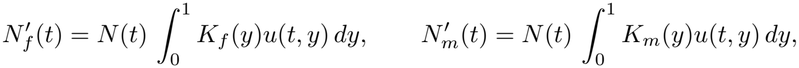

with *N* (*t*) the total gene population. The instantaneous fitness becomes

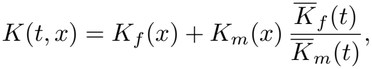

where 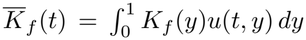 and 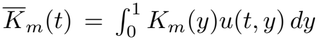 are the mean production rates of female and male gametes in the whole population at time *t*. The mean Malthusian fitness is then

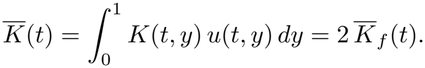

Subtracting 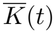 converts absolute growth into relative growth 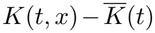 and conserves total genetic mass. This justifies the replicator component of the PDE.

Under Assumption 5, the effect of recurrent mutations with small phenotypic effects can be described by a diffusion approximation. More precisely, if mutations occur at rate *U* and the phenotypic change caused by a single mutation is centered and has variance 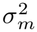, the corresponding diffusion coefficient is 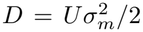 (see Appendix A in Hamel et al. (2020)). Thus, *D* combines the mutation rate and the typical squared magnitude of mutational effects. In the following we call *D* the *mutational diffusivity*. The effect of mutational diffusion is determined by the magnitude of *D* relative to the fitness differences 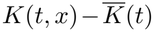 that drive selection. At the population level, this diffusion approximation is represented by the term *D ∂*_*xx*_*u*. Depending on the scale of the mutation (diffusion) parameter *D* compared to selection, the mutational regime will be considered as weak (*D* → 0), intermediate (*D* = *O*(1)), strong (*D* ≫ 1), or infinite (*D* → ∞). In addition, we will also consider cases where there is no mutation (*D* = 0) that allow us to recover classical ESSs results (Charnov et al., 1976; Charnov, 1979, 1982; Lesaffre et al., 2024).

Finally, under Assumptions 1–5, the resulting replicator-mutator PDE governing the evolution of the distribution of trait *x* in the population is

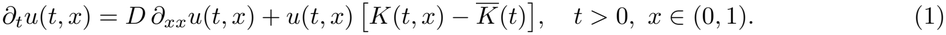

At the boundaries of the trait space, the Neumann conditions

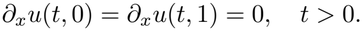

impose zero mutational flux. They conserve genetic mass under the diffusion process while preventing mutations from producing trait values outside the admissible interval [0, 1].

Equation (1) belongs to the classical family of replicator-mutator equations for continuously structured populations (Kimura, 1965; Bürger, 1986; Alfaro and Carles, 2014, 2017; Alfaro and Veruete, 2019). However, the present model has specific features that set it apart from previous models. In particular, unlike in these models, *K* = *K*[*u*](*t, x*) is not a prescribed fitness landscape but a nonlinear, nonlocal functional of the current trait distribution *u*. This dependence arises directly from sexual reproduction, through competition among male gametes for the female gametes produced by the whole population. Thus, although the mutation-selection structure is closely related to replicator-mutator models previously studied mainly for asexual populations, the selective component is here generated endogenously by mating. In contrast with infinitesimal-model formulations (Barton et al., 2017), sexual reproduction therefore does not appear as a separate reproduction operator, but is incorporated into the effective fitness functional *K*[*u*].

### 2.3 Evolutionary stationary states

For any fixed *D >* 0, we denote by *p*_*D*_ a normalized stationary density of (1). Proposition A.1 shows that equation (1) admits a unique positive stationary density *p*_*D*_ for all *D >* 0. Thus, although we do not address the convergence of the time-dependent solution here, any long-time limit, whenever it exists, must coincide with *p*_*D*_. Whenever possible, we derive results for general non-decreasing female fitness functions *K*_*f*_ and non-increasing male fitness functions *K*_*m*_. When general results are not possible to obtain, or when numerically solving the model for stationary distributions, we consider the power-law functions

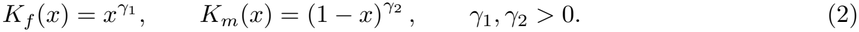

The stationary equation has a useful spectral interpretation. Define

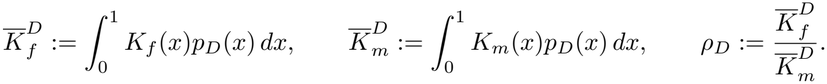

Then *p*_*D*_ satisfies

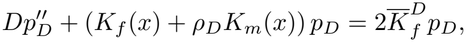

together with

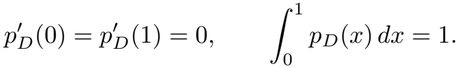

Unlike standard replicator-mutator equations with a prescribed fitness landscape, such as those considered in Alfaro and Carles (2017), the operator in this eigenvalue problem is not fixed in advance. The stationary density *p*_*D*_ determines both its own effective fitness landscape, through *ρ*_*D*_, and the corresponding eigenvalue, through 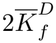 . A stationary state must therefore be a principal eigenfunction while simultaneously satisfying these two consistency conditions. This feedback is the source of the nonlinearity of the stationary problem and makes existence and uniqueness nontrivial.

For *D >* 0, the stationary state *p*_*D*_ is a positive density on [0, 1]. In the singular regimes *D* = 0 and *D* → 0, stationary distributions are instead described as probability measures, denoted by *p*_0_, which may take the form

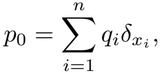

where 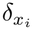 denotes the Dirac probability measure concentrated at the trait value *x*_*i*_, and *q*_*i*_ is the corresponding mass. Across the different mutation regimes, stationary distributions are classified according to the nature of their support and the number and location of their modes. Continuous stationary densities may be uniform, unimodal, or bimodal, with modes located in the interior of (0, 1), at one boundary, or at both boundaries. Altogether, we found eleven qualitatively distinct stationary distributions, summarized in Fig. 7. To name our eleven stationary states without introducing new terminology, we assigned each the name of the commonly considered reproductive system it most closely resembles (hermaphroditism, dioecy, androdioecy, and gynodioecy), adding the suffix “-like” where appropriate (Fig. 7). For example, we use “dioecy” to refer to a stationary distribution concentrated at the two boundaries, corresponding to coexisting pure males and pure females, whereas “dioecy-like” refers to a continuous stationary distribution spanning the entire range [0, 1] and exhibiting two modes, one at each boundary. We also define three additional reproductive systems: male-biased and female-biased, when there is a single mode at one of the boundary, and uniform when there is no mode in the whole range [0, 1]. Trioecy and polygamy are also terms sometimes used to describe reproductive systems, although they are relatively uncommon and sometimes debated. We do not consider them here, as they are not needed to describe the stationary distributions obtained in our model.

Analytical or asymptotic results are obtained in the regimes *D* = 0, *D* → 0, and *D* ≫ 1 (see Appendix A for details). For intermediate values *D* = *O*(1), explicit solutions are generally unavailable. We therefore compute stationary states numerically using the spectral formulation described above (see Appendix A.5 for details). This procedure reduces the nonlinear stationary problem to a sequence of linear principal eigenvalue problems and a scalar consistency equation. It directly targets the stationary state and thus avoids approximating it by integrating equation (1) up to an arbitrarily chosen terminal time.

## 3 Results

### 3.1 Symmetric trade-offs imply symmetric distributions

Assume that male and female fitness functions are symmetric, i.e. *K*_*f*_ (*x*) = *k*(*x*) and *K*_*m*_(*x*) = *k*(1 − *x*), for any continuous functional form of the fitness function *k*. Then the stationary distribution *p* = *p*_*D*_ is symmetric, has a single mode at *x* = 1*/*2 if *k* is strictly concave in (0, 1), or two modes at the boundaries *x* = 0 and *x* = 1 if *k* is strictly convex (see Proposition A.5 in Appendix A.2). This result was previously established (Charnov et al., 1976; Charnov, 1979), but, to the best of our knowledge, only for particular classes of fitness functions.

This general result is true for any *D >* 0, including when fitness functions are power-law *K*_*f*_ (*x*) = *x*^*γ*^, *K*_*m*_(*x*) = (1 − *x*)^*γ*^ with *γ >* 0 (on a side note, the linear case *γ* = 1 is singular and yields *p* ≡ 1). In this particular case of power-law fitness functions, this general result implies that reproductive systems such as gynodioecy and androdioecy, as well as analogous systems characterized by a continuous distribution of sex allocation *x* with one mode in (0, 1) and another at either *x* = 0 or *x* = 1, can arise only when the fitness trade-offs are asymmetric.

Note that the reverse is not necessarily true: the stationary distribution of sex allocation *p* can be symmetric even though the fitness functions *K*_*f*_ and *K*_*m*_ are not. Indeed, the stationary distribution can contain only pure males or females even with asymmetric *k* functions, yet with a balanced sex ratio (Shaw and Mohler, 1953).

### 3.2 Stationary states in the weak mutational diffusion regime (*D* → 0) vs. evolutionarily stable strategies in the no mutation regime (*D* = 0)

In this section, we consider that the trade-offs follow power-law functions *K*_*f*_ (*x*) = *x*^*γ*1^ and *K*_*m*_(*x*) = (1 − *x*)^*γ*2^, *γ*_1_, *γ*_2_ *>* 0.

The stationary states *p*_0_ only show discrete states under the weak mutational diffusion regime (*D* → 0) as they are always composed of one or more Dirac measures (Appendix A.4). It means that at selection-equilibrium, populations are composed of individuals exhibiting a finite number of distinct sex-allocation traits within or at the boundaries of the range (0, 1). This can include situations with pure males, pure females, or hermaphrodites.

In addition, the stationary distributions can be classified into four regions in the parameter space depending on whether *K*_*f*_ and *K*_*m*_ are accelerating (convex, *γ*_*i*_ *>* 1) or decelerating (concave *γ*_*i*_ *<* 1) functions (Fig. 2 and Theorem A.10):

**Figure 2:**
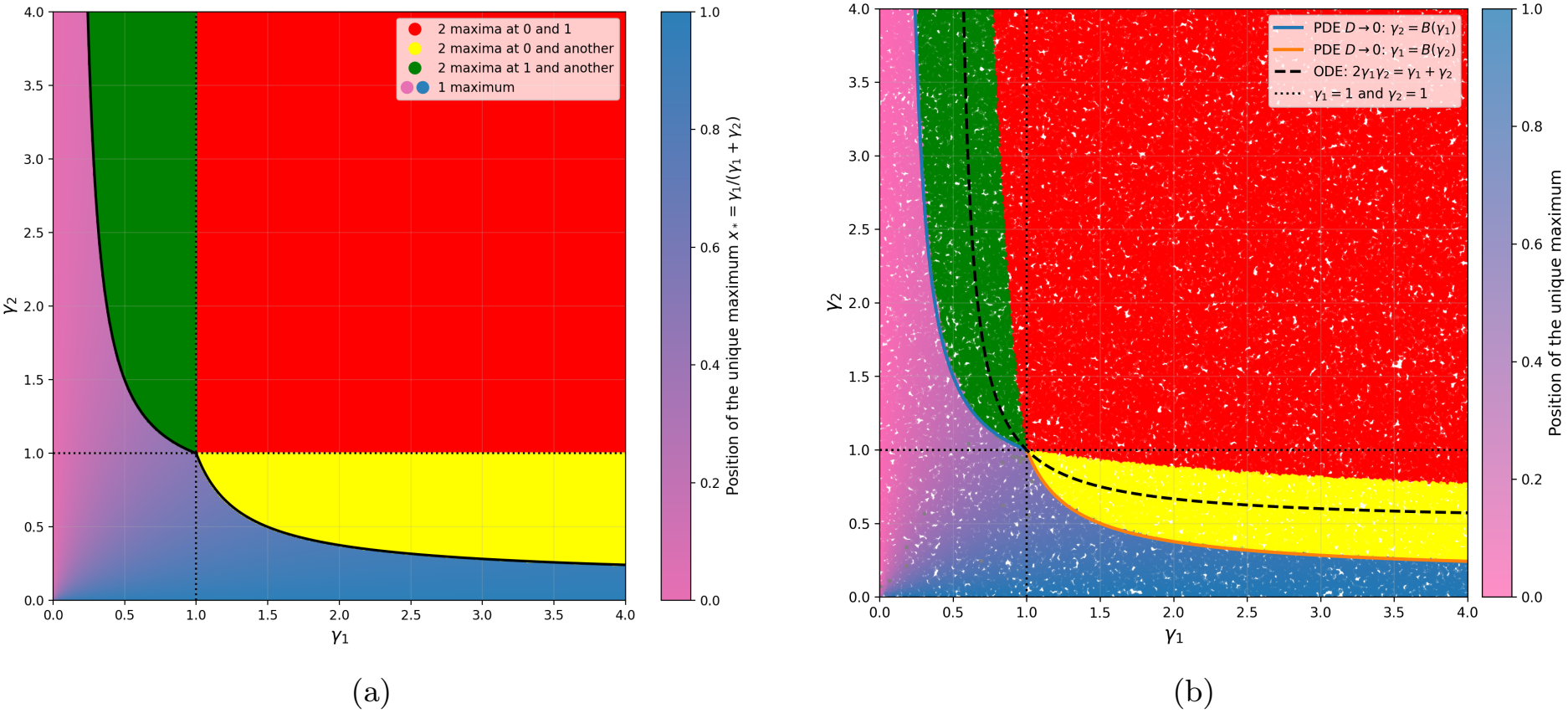
Stationary distributions predicted under the weak mutational diffusion limit of the PDE (1). (a): phase diagram predicted as the exact asymptotic limit when *D* → 0 of (1) (Theorem A.10) in the limit *D* → 0. (b): phase diagram obtained numerically with *D* = 10^*−*10^. The black dashed curves and the thick curves are the one obtained under the no mutation regime *D* = 0, respectively corresponding to the results by Lesaffre et al. (2024) and Charnov (1979) (Appendices C.1 and C.2). Color refers to the different stationary states of sex allocation (position of the Dirac measures): the pink-to-blue gradient corresponds to a unique interior trait (hermaphroditism), red to two traits at 0 and 1, yellow to two traits at 0 and one interior point (androdioecy), and green to two maxima at 1 and one interior point (gynodioecy).

1. If *γ*_1_ *<* 1 and *γ*_2_ *<* 1, or *γ*_1_ *<* 1 *< γ*_2_ ≤ *γ*_1_ (2^1*/γ*1^ – 1), or *γ*_2_ *<* 1 *< γ*_1_ ≤ *γ*_2_ (2^1*/γ* 2^ – 1):
  → Hermaphroditism: *p*_0_ = *δ*_*x*_, i.e hermaphrodites with sex allocation 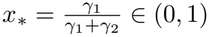;
2. If *γ*_1_ *>* 1 and *γ*_2_ *>* 1:
  → Dioecy: 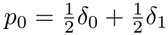, i.e. pure males and females with a balanced sex ratio;
3. If *γ*_1_ *<* 1 and 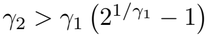:
  → Gynodioecy: *p*_0_ = *q*_*s*_*δ*_*s*_ + (1 − *q*_*s*_)*δ*_1_, i.e. hermaphrodites with sex allocation *s* ∈ (0, 1) and females with respective frequencies *q*_*s*_ ∈ (0, 1) and 1 − *q*_*s*_ ∈ (0, 1);
4. If *γ*_2_ *<* 1 and *γ*_1_ *> γ*_2_ (2^1*/γ* 2^ – 1):
  → Androdioecy: *p*_0_ = (1 − *q*_1*−t*_)*δ*_0_ + *q*_1*−t*_*δ*_1*−t*_, i.e. hermaphrodites with sex allocation 1 − *t* ∈ (0, 1) and males with respective frequencies *q*_1*−t*_ ∈ (0, 1) and 1 − *q*_1*−t*_ ∈ (0, 1).

The explicit values of the allocation to female function in hermaphrodites *s* and *q*_*s*_, as well as the frequency 1 − *t* and *q*_1*−t*_ of hermaphrodites in gynodioecious or androdioecious populations are given in Theorem A.10.

The stationary distributions obtained from numerical estimation for a very small but not null mutational diffusivity (*D* = 10^*−*10^) show few differences with the asymptotic solutions when *D* → 0 (compare Fig. 2a and Fig. 2b). More specifically, the regions delimiting hermaphroditism are very similar (pink to blue region delimited by the thick black curve). The region delimiting dioecy slightly differs because of numerical approximations when determining if the mode is exactly at the boundaries.

It was intriguing that under the weak mutational diffusion regime *D* → 0, we recovered the results from Charnov (1979), and only partially the results from Lesaffre et al. (2024). The latter indeed found the hermaphroditism region to be larger and delimited by 2*γ*_1_*γ*_2_ = *γ*_1_ + *γ*_2_ (the black dashed curve in Fig. 2). We therefore analyzed our PDE model (1) in a situation analogous to theirs by setting *D* = 0 and studying the dynamics of an initial distribution supported on only two traits: the resident 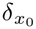 and an initially rare mutant 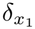 . We found the same results as Lesaffre et al. (2024) when assuming that the mutant was close to the resident, and the same results as Charnov (1979) when relaxing this assumption, *i*.*e*. when the mutant can appear anywhere in the phenotypic space [0, 1]. This includes, in particular, the appearance of pure males or females in a hermaphroditic resident population (see Appendix C). Our PDE model is thus general enough to recover classical results based either on an explicit diploid genetic model (Charnov, 1979) or on the adaptive dynamics approach (Lesaffre et al., 2024).

Yet, in our PDE model, mutations are introduced through the diffusion term in the PDE (1), i.e. assuming that mutations act locally in trait space. Since 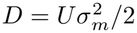, the limit *D* → 0 may result from a decreasing mutation rate *U*, from mutational effects of decreasing variance 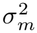, or from both. Thus, depending on how this limit is approached, mutations may become rarer, have increasingly small effect, or both. Why, then, do we recover the global invasibility criterion of Charnov (1979) rather than the local criterion of Lesaffre et al. (2024) based on nearby mutants?

Our asymptotic results are obtained by considering the limits in the following order. For each fixed *D >* 0, we first determine the stationary distribution *p*_*D*_, formally corresponding to the limit *t* → +∞. We then study the limit of *p*_*D*_ as *D* → 0. Hence, even though mutations act locally, the first step allows time to be large enough such that the distribution of sex allocation covers the whole range [0, 1], including the boundary phenotypes corresponding to pure males and females. The limit is therefore governed by global invasibility, as in Charnov (1979).

### 3.3 Gynodioecy- and androdioecy-like evolutionary stationary states are absent in the strong-mutation regime (*D* ≫ 1)

Under the strong mutational diffusion regime (*D* ≫ 1), assuming that trade-offs follow power-law functions (2) with *γ*_1_≠ *γ*_2_, we found four possible stationary states depending on the convexity of *K*_*f*_ and *K*_*m*_, all stationary states being continuous and covering the whole range (0, 1) (Fig. 7). These stationary states are characterized by the number and localization of their modes (Appendix A.3):

1. If *γ*_1_ *<* 1 and *γ*_2_ *<* 1: Hermaphroditism-like, *i*.*e*. a unique mode at *x*_0_ within (0, 1);
2. If *γ*_1_ *>* 1 and *γ*_2_ *>* 1: Dioecy-like, *i*.*e*. two modes at boundaries *x* = 0 and *x* = 1;
3. If *γ*_1_ *<* 1 and *γ*_2_ *>* 1: Male-biased, *i*.*e*. a unique mode at *x* = 0;
4. If *γ*_2_ *<* 1 and *γ*_1_ *>* 1: Female-biased, *i*.*e*. a unique mode at *x* = 1;

Note that gynodioecy-like and androdioecy-like stationary states are absent, and that male-biased and female-biased distributions are original stationary states with no counterpart when *D* → 0 or *D* = 0. An illustration of male-biased stationary state is shown in Fig. 5a for a mutational diffusivity *D* = 10^*−*1^. When trade-offs *K*_*f*_ and *K*_*m*_ follow power-law functions, we can compare the parameter space of the stationary states between the weak (*D* → 0) vs. strong (*D* ≫ 1) mutation regime (Fig. 2a and 3f vs. Fig. 3a). First, with strong mutation, hermaphroditism-like occurs in a much narrower region than hermaphroditism: hermaphroditism-like is limited to the unit square (0, 1) × (0, 1) whereas hermaphroditism can occur even for large values of *γ*_1_ or *γ*_2_. Hence, strong mutational diffusion tends to eliminate interior modes and push modes to the right or left boundaries. Second, dioecy-like and dioecy occur in the same parameter space: the square (1, ∞) ×(1, ∞). Hence, even if strong mutational diffusion tends to eliminate interior modes, it does not increase the region where two modes at both boundaries occur. Finally, male-biased distribution covers the region where either hermaphroditism or gynodioecy occur, and female-biased distribution covers the region where either hermaphroditism or androdioecy occur. Hence, gynodioecy and androdioecy, or even gynodioecy-like and androdioecy-like (see next section) stationary states are eliminated by strong mutation.

**Figure 3:**
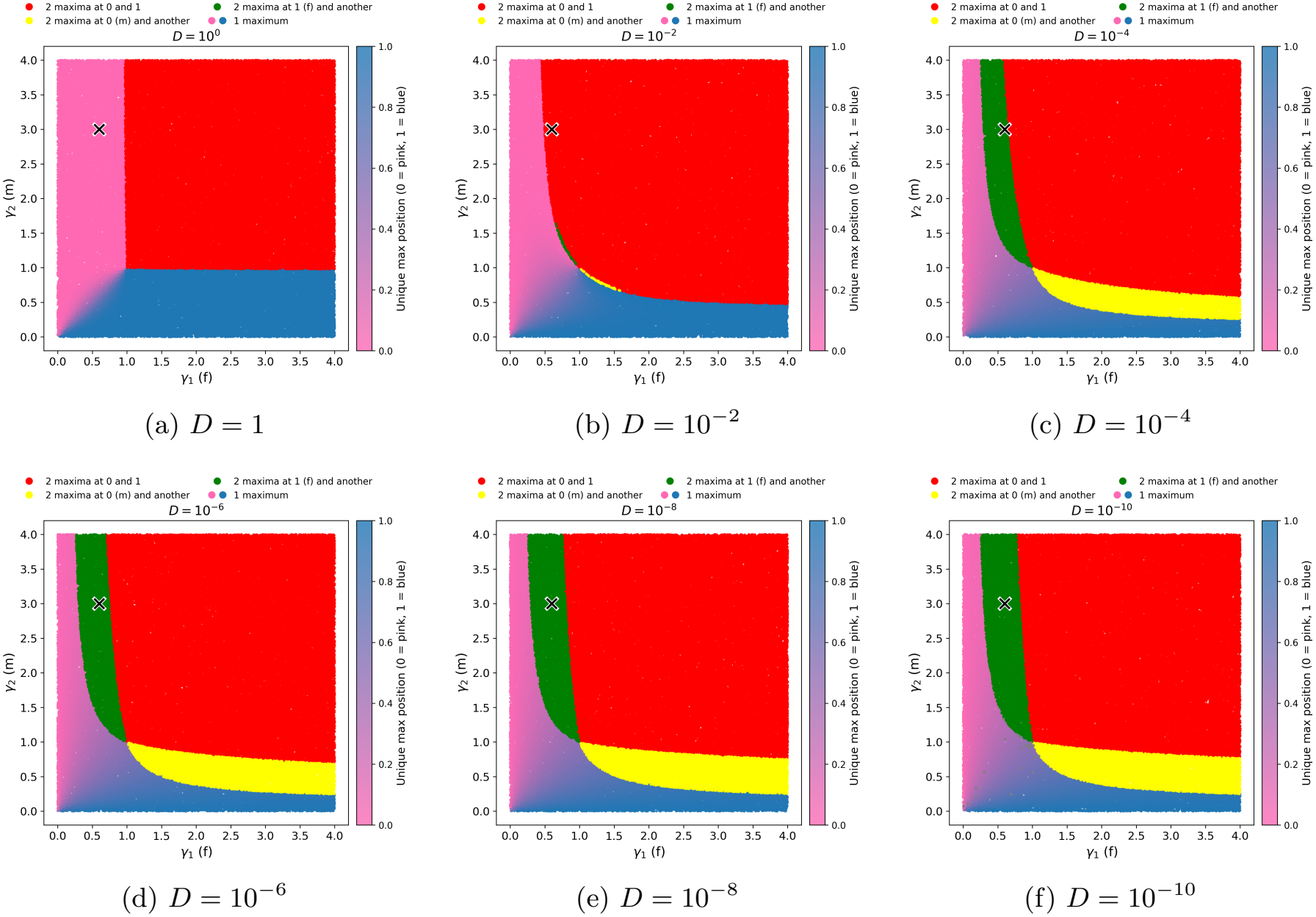
Stationary distributions *p*_*γ* 1,*γ*2_ for different values of *D* (see Fig. 7 for terminology). Red: stationary distributions with two modes at 0 and 1 (‘balanced’ or ‘unbalanced dioecy-like’); Yellow: two modes, one at 0 (male) and the other at an interior point (‘androdioecy-like’); Green: two modes, one at 1 (female) and the other at an interior point (‘gynodioecy-like’): Pink-blue gradient: correspond to stationary distributions with a unique mode, whose position varies continuously from 0 (pink) to 1 (blue) (in that order: ‘male biased’, ‘hermaphroditism-like’, ‘female-biased’). The black cross marks the parameter pair (*γ*_1_, *γ*_2_) = (0.6, 3) corresponding to Fig. 5.

On a side and final note, in an even stronger mutation regime than *D* ≫ 1, *i*.*e*. in the infinite mutation regime *D* → +∞, we show in Proposition A.6 that the unique stationary state *p*_*D*_ converges to the uniform density *p*_*∞*_ ≡ 1 for any trade-offs function *K*_*f*_ and *K*_*m*_, including power-law functions. This is not surprising as mutation is the unique process governing the evolution of the population (selection is negligible under this regime).

### 3.4 Intermediate mutation regime (*D* = *O*(1)) shows that mutation alone can drive reproductive systems shifts

We showed above that the distribution of sex allocation can greatly differ under different asymptotic mutation regimes (compare *D* → 0 and *D* ≫ 1). We now explore how the value of *D* affects the distribution of sex allocation for any intermediate values. General explicit analytical results were out of reach for any *D*, we thus relied on numerical estimations of stationary distributions assuming that trade-offs followed power-law functions *K*_*f*_ (*x*) = *x*^*γ* 1^ and *K*_*m*_(*x*) = (1 − *x*)^*γ* 2^, *γ*_1_, *γ*_2_ *>* 0.

We found seven possible stationary distributions when *D* was intermediate, represented on Fig. 3 by regions of different colors: balanced or unbalanced dioecy-like (two modes at both boundaries - red); gynodioecy-like (one mode at boundary *x* = 1, one mode within (0, 1) - green); androdioecy-like (one mode at boundary *x* = 0, one mode within (0, 1) - yellow); male-biased (one mode at boundary *x* = 0 - pink); female-biased (one mode at boundary *x* = 1 - blue); hermaphroditism-like (one mode within (0, 1) - shades of purple); uniform (no modes). Hence, compared to the stationary sex allocation distribution when mutation is strong *D* ≫ 1, two other reproductive systems were found: androdioecy-like, gynodioecy-like.

Fig. 3 shows how the regions of parameter space (*γ*_1_, *γ*_2_) corresponding to each of these possible stationary sex-allocation distributions vary with the value of *D*. As expected, the diagrams for very small (*D* = 10^*−*10^) or large (*D* = 1) mutational diffusivity are very close to asymptotic limits (respectively *D* → 0 and *D* ≫ 1). Fig. 3 also shows that androdioecy-like and gynodioecy-like distributions occur for narrower parameters space as *D* increases. This is due to an expansion of the parameter space where dioecy-like distributions occur: for intermediate *D* values, modes at both boundaries can occur even though one trade-off function is concave hence decelerating, which is illustrated on Fig. 4c where *D* = 10^*−*4^ and *γ*_1_ = 0.8. This is a specificity of intermediate values as dioecy or dioecy-like distributions were limited to cases where both trade-offs functions were accelerating under weak (*D* → 0) or strong (*D* ≫ 1) mutation regimes.

**Figure 4:**
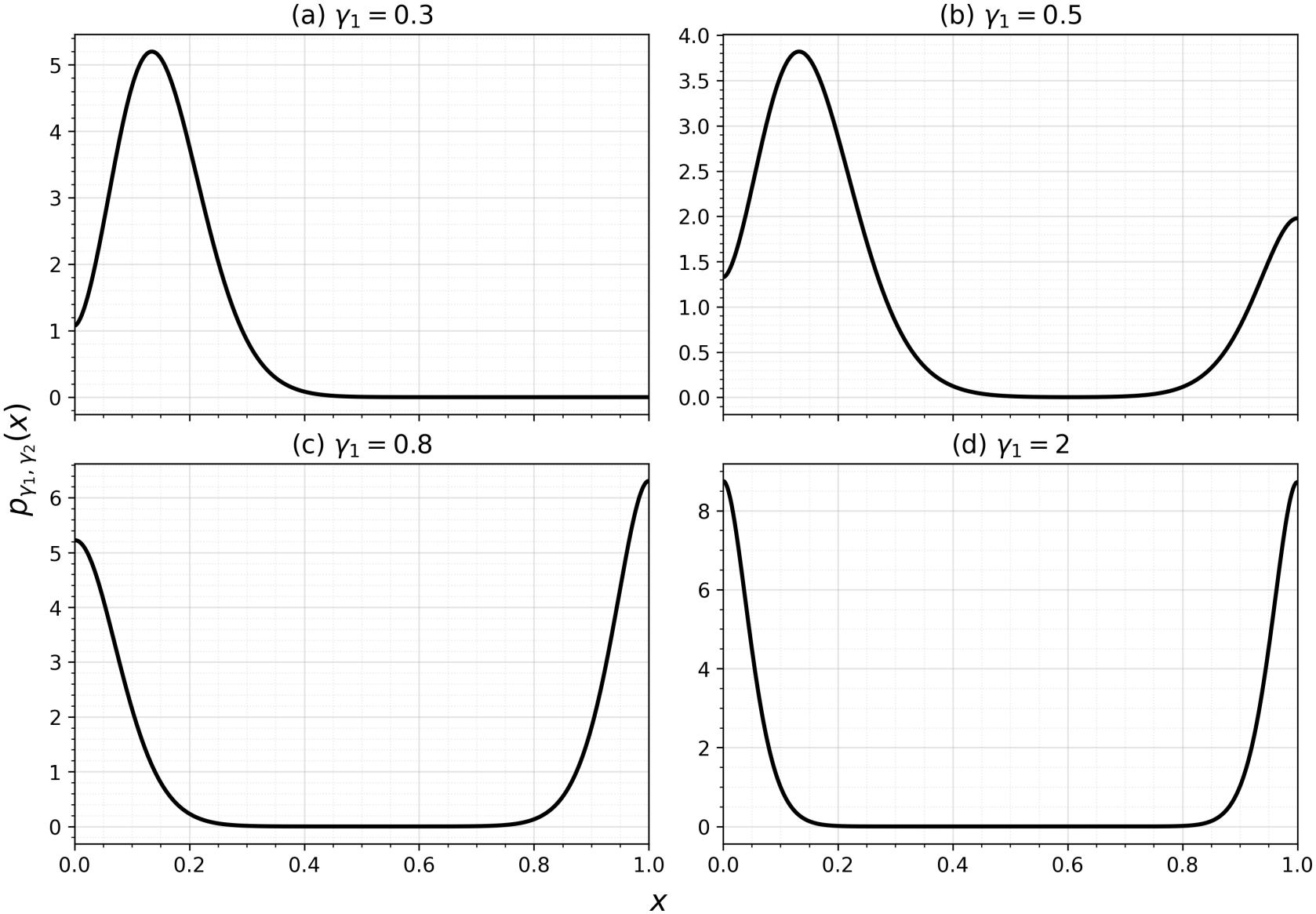
Stationary distributions *p*_*γ* 1,*γ*2_ for *D* = 10^*−*4^, *γ*_2_ = 2 and increasing values of *γ*_1_. (a): hermaphroditism-like; (b): gynodioecy-like; (c): unbalanced dioecy-like ; (d): balanced dioecy-like.

Strikingly, the variation of the parameter space where dioecy-like occurs is not monotone with respect to *D*: it first expands eliminating gynodioecy-like and androdioecy-like sex allocation (compare Fig. 3f to Fig. 3b) and then shrinks back because of the expansion of male-biased and female-biased sex allocation (compare Fig. 3b with Fig. 3a). As an illustration, Fig. 5 shows how stationary distributions vary with *D* for *γ*_1_ = 0.6 and *γ*_2_ = 3 (corresponding to the black cross on Fig. 3). The stationary sex allocation distribution goes from gynodioecy-like (green on Fig. 3), to dioecy-like (red) and finally to male-biased (pink) as *D* increases. Hence, for given trade-offs function and associated parameters *γ*_1_ and *γ*_2_, mutation alone can drive a shift in reproductive systems.

**Figure 5:**
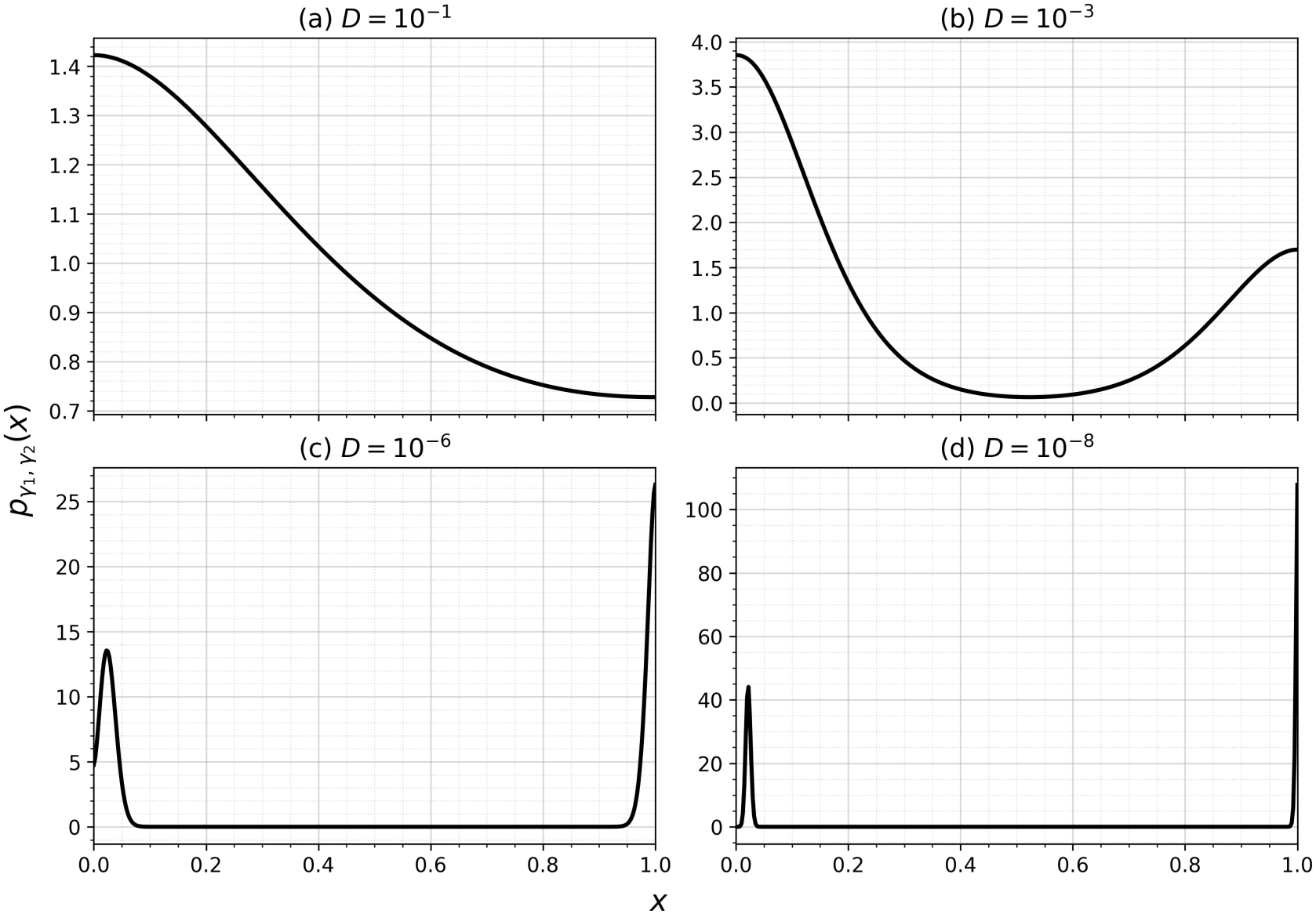
Stationary distributions *p*_*γ* 1,*γ*2_ for *γ*_1_ = 0.6 and *γ*_2_ = 3, for decreasing values of the mutational diffusivity *D*. (a): *D* = 10^*−*1^, male-biased; (b): *D* = 10^*−*3^, unbalanced dioecy-like; (c) *D* = 10^*−*6^ and (d) *D* = 10^*−*8^, gynodioecy-like.

How can mutation drive a reproductive system shift on its own? This is because the fitness *K*[*u*](*t, x*) of a phenotype *x* is frequency-dependent and depends on the distribution of sex allocation in the whole population *u*. As mutation affects the distribution of traits *u*(*t, x*), it consequently affects *K*[*u*](*t, x*) and thus the stationary distribution. In addition, when male and female fitness trade-offs functions are asymmetric, modes of the distribution can occur close to the boundaries. As boundaries are reflective, the rate at which mutants are introduced in the populations is not symmetrical on both sides of the mode, generating an overproduction of male or female gametes, thus affecting the location of the modes of the stationary states.

This can be illustrated by the shift from gynodioecy-like to dioecy-like and male-biased sex allocation distribution when *D* increases (Fig. 5). When mutational diffusivity is very small (*D* = 10^*−*8^), selection dominates mutational diffusion which makes modes concentrated around the optimal value. Increasing mutational diffusion relative to selection tends to send traits further away from the optimum (compare Figs. 5c and 5d). Because the interior mode is close to the boundary *x* = 0 and because the boundary is reflective, mutants tend to be more frequent on the right of the peak than on the left, i.e. mutants coming from this optimum tend to produce more female gametes. This creates an asymmetry favoring individuals producing more male gametes pushing the optimum to the boundary *x* = 0, thus producing a shift from gynodioecy-like to dioecy-like reproductive systems. At the same time, the proportion of pure females decreases: females are more than twice more frequent than hermaphrodites at the mode close to *x* = 0 when *D* = 10^*−*8^, but as *D* increases females proportion decreases until being less frequent than males (*D* = 10^*−*3^). When *D* is large, the mode at *x* = 1 eventually disappears generating a shift from dioecy-like to male-biased distributions. Note that these two successive shifts eliminating pure females and favoring pure males as *D* increases occur in the region where *γ*_1_ *<* 1 and *γ*_2_ *>* 1, i.e. where the male trade-off is accelerating and the female trade-off is decelerating. The opposite shifts occur in the reverse situation.

These shifts however come at a cost as the mean fitness tends to decrease when the mutational diffusivity increases, due to the recurrent introduction of deleterious mutants, thus generating a mutation load (Fig. 6). Indeed, when *D* is small mean fitness spans from 1 (dioecy-like) to 2 (male-biased and female-biased), while when *D* is large it spans from 0.5 (dioecy-like and female-biased) to 2 (male-biased). Hence, mutation load at the genes underlying sex allocation can be associated with reproductive system shifts.

**Figure 6:**
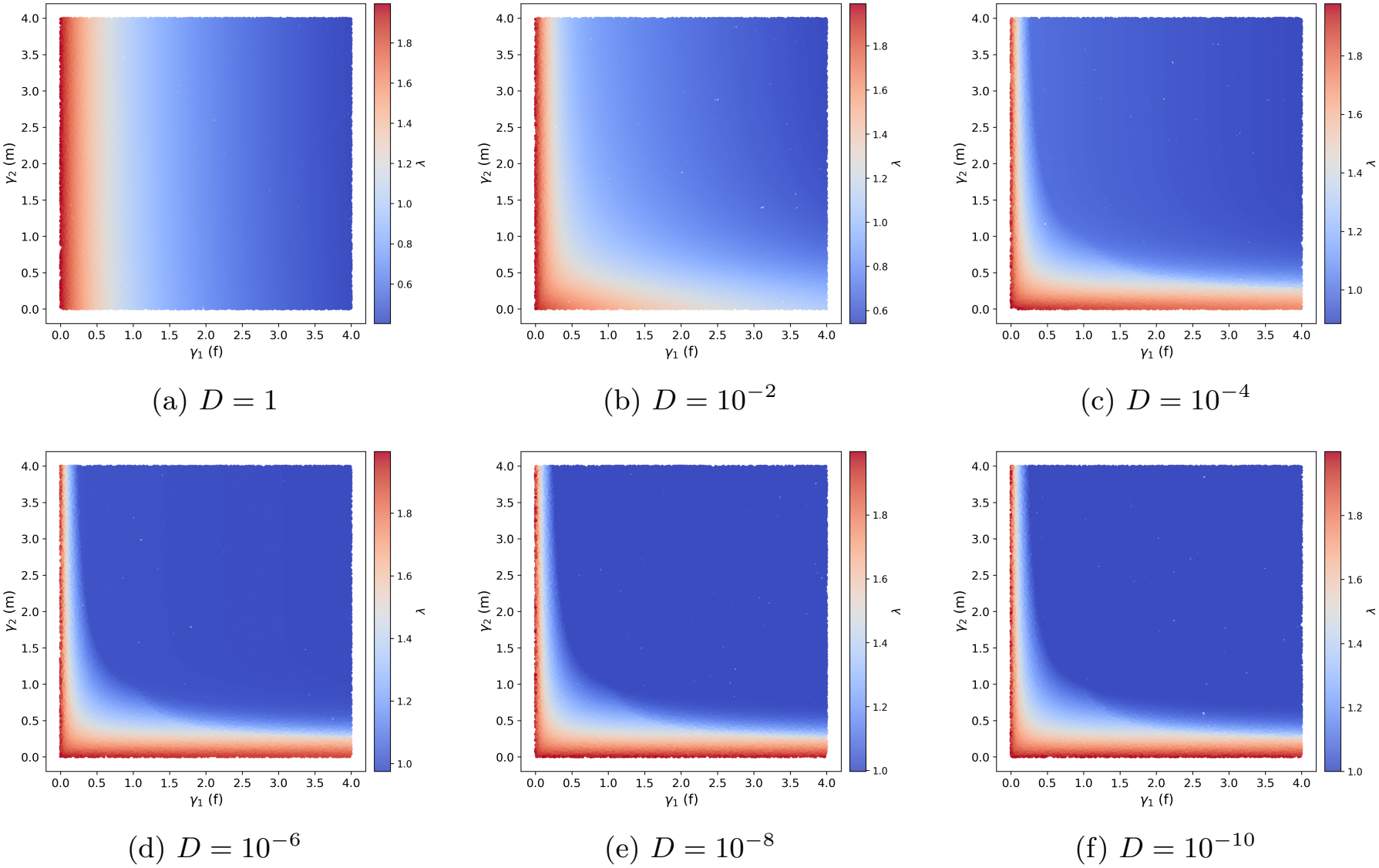
Mean fitness 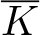 at stationary distribution for different values of *D* (see Fig. 3 for the corresponding stationary distributions).

## 4 Discussion

### 4.1 Which reproductive systems are evolutionarily stable?

Reproductive systems are generally classified according to discrete sexual phenotypes based on whether individuals produce male or female gametes. Individuals are thus classified as males, females, or hermaph-rodites. Reproductive systems are then defined according to which sexual phenotypes coexist within a population. Four reproductive systems are most commonly described : dioecy, hermaphroditism, gynodioecy and androdioecy (or their equivalent in haplodiplontic or haplontic species). These four reproductive systems are indeed found as evolutionarily stable under the simplest models where only competition between male gametes and allocation trade-offs are assumed (Charnov et al., 1976; Lesaffre et al., 2024). This correspondence between the prediction of the simplest models with the highly prevalent four main reproductive systems showed a large success in evolutionary biology for explaining the diversity of reproductive systems, as well as their evolutionary transitions. This success further suggested that any other observed reproductive systems would either: 1) be evolutionary unstable and transitory to one of the four stable states (Charnov et al., 1976; Maurice and Fleming, 1995; Perry et al., 2012; Adams et al., 2025); 2) need some extra ecological or developmental mechanisms playing an important role, but possibly under restricted parameter regions, such as pollen limitation (Maurice and Fleming, 1995) or maternal effect (Adams et al., 2025); 3) show that sex phenotyping was biased or wrongly reported (see for example the controversy about the functional role of hermaphrodites in *Phillyrea angustifolia* Francq et al. 2023).

Yet, when looking at sex phenotypes in a quantitative way, data often show a continuous rather than a discrete distribution of, at least partly heritable, phenotypic variation. It is so common in plants, animals or mosses, that many other categories have been introduced in the literature such as trioecy, subdioecy or polygamy when males, females and hermaphrodites co-occur depending on the proportion of hermaphroditic individuals in the population, as well as the degree of specialization to one sex. Hermaphroditic species can also be described in finer categories depending on the types of organs (e.g. flowers) individuals produce at a given time, possibly reflecting the resource allocation to the production of one type of gametes or the others. In flowering plants for example, populations can be composed of individuals with perfect flowers in which both types of gametes are produced simultaneously (true hermaphroditism), or individuals with specialized flowers producing either male or female gametes separated in space (monoecy) or time (dichogamy), or individuals with male and bisexual flowers (andromonoecy), and many other categories. A similar categorization is used in mosses also possibly reflecting different sex allocation between individuals (dos Santos et al., 2024; Dos Santos et al., 2026).

In the current paper, we showed that many of the reproductive systems described in natural populations can be recovered with a single model with minimal assumptions (Fig. 7), only by adding small recurrent mutations to the well-established mechanisms in the fundamental models of sex allocation theory (Charnov et al., 1976). This includes the four main discrete distributions commonly used to describe reproductive systems (hermaphroditism, dioecy, androdioecy and gynodioecy), but also others like subdioecy or polygamy that can correspond to different stationary states predicted by our model, such as dioecy-like, gynodioecy-like, androdioecy-like or hermaphroditism-like (Fig. 7). Our model thus shows that intermediate reproductive system such as polygamy can indeed be evolutionarily stable, a question which is still debated in the literature (e.g. Saumitou-Laprade et al. 2018; Godin 2022). Depending on the mutational diffusivity and the form of the sex allocation trade-offs we eventually found eleven qualitatively different stationary distributions that could be used to more accurately describe the reproductive systems found in nature. This includes original reproductive systems with no equivalent to our knowledge in the literature: male-biased, female-biased or uniform.

**Figure 7:**
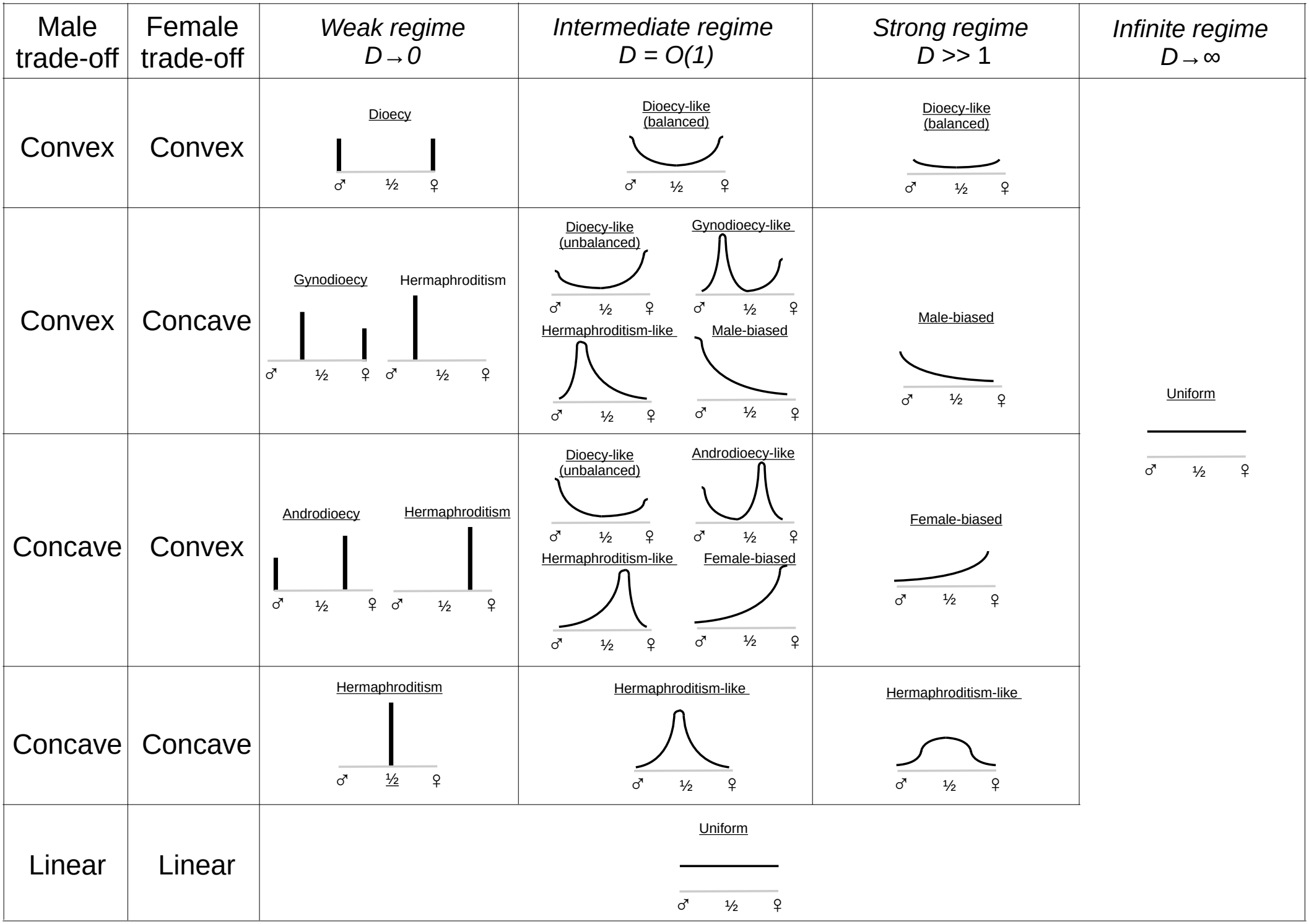
Schematic representation of the eleven stationary distributions of sex allocation (reproductive systems) found in our model, depending on the form of the trade-off functions and the mutation regime. The stationary distributions can be composed of a finite number of traits (Dirac measures) when *D* → 0, otherwise they are continuous and cover the whole range (0, 1). The classical terminology ‘dioecy’, ‘androdioecy’, ‘gynodioecy’, ‘hermaphroditism’ was kept when the stationary distributions are discrete. An analogous terminology is introduced when the distributions are continuous based on their resemblance with the classical terminology and the number and localization of the distribution modes: ‘dioecy-like (balanced)’ or ‘dioecy-like (unbalanced)’ (two symmetric or asymmetric modes at the boundaries), ‘androdioecy-like’ (one mode at the male boundary, another within (0, 1)), ‘gynodioecy-like’ (one mode at the female boundary, another within (0, 1)), ‘hermaphroditism-like’ (one mode within (0, 1)). A new terminology is introduced for other cases: ‘male-biased’ (one mode at the male boundary), ‘female-biased’ (one mode at the female boundary), ‘uniform’ (no modes).

There is yet at least one reproductive system which is not predicted by our model: trioecy where three modes are observed, one at each boundary corresponding to pure males and females and one corresponding to hermaphrodites (e.g. Fleming et al. 1998). The first possibility is that our model lacks one key element to be able to predict trioecy, such as pollen limitation as suggested by Maurice and Fleming (1995). However, even if we include pollen limitation in our model, it is not guaranteed that stationary distributions with three modes could emerge from it rather than the flat distribution of hermaphrodites between males and females such as in the case of the ‘dioecy-like’ situations (Fig. 7). The second possibility is that what is called trioecy in natural populations actually shows such a flat continuous distribution of hermaphrodites as in ‘dioecy-like’, and that the third mode is artificially produced by aggregating all hermaphroditic individuals within a single class.

Finally, there is another possible discrepancy between the predictions of our model and observed distribution of sex phenotypes in populations that would need further investigations. In *Veronica subalpina* (Fig. 1a), there are two interior modes, a situation not predicted by our model. In our model, when the stationary distribution has two modes, at least one of them is necessarily at one of the boundaries. Here again it might suggest that some key mechanisms are absent in our model. Another possibility is that the proxy used to estimate the resource allocation, here the fruit-set, does not directly reflect resource allocation. How to correctly estimate resource allocation to sexes from experiments or natural populations is a long-standing debate in the literature with no consensus to our knowledge.

### 4.2 Sources of phenotypic variations

We hypothesized that the stationary distributions of sexual phenotype variations were only due to a balance between the rate of small recurrent mutations, and the strength of selection mediated by reproductive trade-offs. There are yet several other documented sources of sexual phenotypic variations, the relative importance of which is debated for long.

We neglected plausible complex interactions with the environment, supposing that environment was constant. It is well-known that in many species sex allocation is for instance mediated by temperature (e.g. in cucumber Lai et al. 2018, or nematodes Yoshida et al. 2024). We also supposed that there was no plastic response to the social environment of individuals whereas some studies suggest it can occur when one sex is rare, or during a colonization process (Adams et al., 2025; Kostur and Wadgymar, 2023).

We also neglected developmental noise. For instance, labile sex expression in flowering plants, where individuals of one sex can produce flowers of the other sex, has been documented in dioecious species whether sex is determined with sex chromosomes or not (e.g. Käfer et al. 2022). More generally, developmental noise (also coined sex inconstancy, leaky dioecy, among other terms) can be a cause of non-heritable sexual phenotypic variation that can partly explain the observed distributions. Our assumption that mutation is the only source of phenotypic variation that can explain the observed sexual phenotypic distribution is thus a simplification. However, several studies (e.g. Delph and Lloyd 1991; Mazer et al. 1999; Saumitou-Laprade et al. 2018) compared sexual phenotypes among non-sibs, sibs, clones, or within individuals across time, or estimated the heritability of traits involved in the expression of sex and the production of gametes. They clearly showed that genetic factors generally play substantial roles. Experimental evolution studies corroborated this claim by showing that sex allocation rapidly and strongly responded to artificial selection (Mazer et al., 1999; Gerchen et al., 2026). Hence, our assumption that mutation is a major mechanism underlying the distribution of sex phenotypic variation seems reasonable.

Our PDE model is general enough that it could be adapted to include other sources of variation, including: A heterogeneous environment in time or space that could affect the sexual expression of a given genotype through a given reaction norm; Plastic response to the social environment, i.e. as a feedback of the distribution of sex allocation in the population at a given time; Developmental noise by decoupling the trait which is inherited and the trait expressed. As selection is frequency-dependent in our model, it is difficult to predict how these additional sources of phenotypic variation would affect the evolutionary stationary states. That justifies further studies using our theoretical framework.

### 4.3 Recurrent mutations affect reproductive systems evolution because of an interaction with frequency-dependent selection

One of our main results is that mutation alone can drive reproductive system shifts, as illustrated in Fig. 5. For a given trade-off and associated parameters, the resulting stationary distribution and reproductive systems shift from gynodioecy, to dioecy-like and then to male-biased distribution when the mutation regime respectively shifts from weak, to intermediate and strong regime. Such a shifting of reproductive systems through mutation alone is a consequence of frequency-dependent selection which is inherent to the evolution of sex allocation when there is competition between male gametes for the access of female gametes. Recurrent mutations constantly introduce a flow of phenotypic variants which affects the distribution of sexual phenotypes in the population. In short, our model shows that the recurrent flow of maladapted individuals can affect the evolutionarily stable state. A similar situation was found where a species’ range was affected by the constant flow of maladapted individuals by migration (Kirkpatrick and Barton, 1997).

This flow of variants by mutation can be asymmetric because of the boundaries which define pure males and pure females. This asymmetry has two striking consequences when the mutational diffusivity increases. First, modes within the range (0, 1) disappear and are pushed to the boundaries then decreasing the parameter space where hermaphroditism is evolutionarily stable. Second, as the mutational diffusivity continues to increase, the mode at the boundary in gynodioecy-like or androdioecy-like reproductive system disappears and the interior mode is pushed to the other boundary, finally giving male-or female-biased distributions. These results illustrate how the interaction between mutation and frequency-dependent selection affects the evolutionarily stable distribution of sexual phenotype in counter-intuitive ways. For instance, when trade-offs are convex and concave, our model shows that dioecy-like reproductive system with unbalanced sex ratio is a possible evolutionarily stable state, whereas without mutation only gynodioecy or androdioecy is expected (Charnov et al., 1976; Lesaffre et al., 2024).

A question raised by these results is whether mutational diffusivity varies sufficiently among species or clades to contribute to shifts in reproductive systems, or at least whether they can be sufficiently high to significantly affect the distribution of sex allocation observed within a population. This question is difficult to address directly, but several lines of evidence suggest that intermediate or high mutational diffusivity regimes may be biologically plausible. First, as noted above, many species exhibit substantial heritable variation in traits involved in sex allocation. Second, the rapid response to artificial selection observed in experimental evolution studies suggests, non-exclusively, either that substantial standing variation is present in the ancestral population or that mutation and recombination can rapidly generate phenotypic variants even from a small number of individuals at the onset of the experiment. Finally, recent genomic studies have shown that the genetic architecture underlying sex-allocation phenotypes involves a large number of genes with variable effects (Gerchen et al., 2026). One might therefore expect the potential mutational diffusivity to increase with the number of genes contributing to the phenotype.

Finally, selection on sexual traits is generally expected to be strong which seems contradictory with their large heritable variation. It is thus difficult to explain how large genetic variation is maintained, as already noted by Mazer et al. (1999). Several explanations have been proposed such as the existence of antagonistic pleiotropic effects of genes and mutations involved in either male or female phenotypic expression. Our model provides a simpler explanation: substantial genetic variation can be maintained at mutation-selection balance when mutational diffusivity is sufficiently large relative to the selective differences generated by sex-allocation trade-offs. Yet, our model implicitly assumes a simple genetic architecture underlying sex allocation traits analogous to quantitative genetics: a very large number of independent genes with small effects. The validity of this approximation remains to be assessed. In our model, *D* is not the mutation rate itself, but the mutational diffusivity 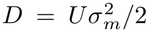, which combines the mutation rate *U* and the variance 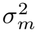 of mutational effects. Its influence must therefore be interpreted relative to the scale of the fitness differences generated by the trade-offs. In the power-law family considered here, the fitness functions are normalized, so increasing *D* increases mutational spreading relative to this fixed fitness scale, producing progressively flatter stationary distributions, as illustrated in Fig. 5. Similarly, the closer the trade-offs are to being linear (*γ*_1_ ≈ 1, *γ*_2_ ≈ 1), the more uniform and flatter the stationary distributions are expected to be, because selection is weaker.

### 4.4 A single PDE model that encompasses and goes beyond previous models

We showed that our model can recover the results obtained previously in phenotypic models (Charnov et al., 1976), population genetics models (Charnov, 1979) and adaptive dynamics models (Lesaffre et al., 2024). We especially show that results that can differ between models are due to different assumptions relatively to how mutations are considered, *i*.*e*. these models can be seen as particular cases of our PDE model.

Previous phenotypic (Charnov et al., 1976), population genetic (Charnov, 1979) and adaptive dynamics models (Lesaffre et al., 2024) consider the invasion of a resident population at equilibrium by a single rare mutant. Such a procedure for searching ESSs makes implicit assumption about the relative scaling of mutation rates, mutation size, and selection intensity. It is not clear whether we can recover the same scaling limits from our PDE model, or if the modelling approaches are structurally different.

Our results nevertheless show that the order in which the limits are considered is crucial. We first determine the stationary distribution *p*_*D*_ for each fixed *D >* 0, formally corresponding to the long-time limit *t* → +∞, and then investigate the regimes *D* → 0, *D* ≫ 1 or *D* = *O*(1). This order of limits explains why the regime *D* → 0 recovers the global invasibility criterion of Charnov (1979), rather than the local criterion of Lesaffre et al. (2024): time is large enough in our model such that even though mutations have infinitesimal effects, the whole range [0, 1] is covered.

This raises the question of the range of validity of our PDE model. Under which conditions on the parameters is our model approximately correct? Answering this question would need to study a stochastic individual-based model where each individual is characterized by a continuous trait *x*, where sexual reproduction is explicitly modeled as the fertilization between male and female gametes under a given mating model, where there is a given genotype-phenotype map characterizing the form of the trade-offs, and for a given mutation kernel and an underlying genetic architecture. Studying to which limit would this microscopic stochastic model converge is necessary to fully assess whether or not our model is a good approximation, and if so under which parameter scaling.

## Data availability

All data and scripts will be published after embargo on a public repository.

## Acknowledgments

SB especially thanks Philippe Vernet and Pierre Saumitou-Laprade for inspiring scientific exchanges for many years, as well as Isabelle De Cauwer and Thomas Lesaffre for helpful discussions, Cassien Milesi Billiard for collecting data points from the figure in Delph and Lloyd (1991), and Sandrine Maurice for her invaluable informatics archaeological work in retrieving and sharing data on *Silene acaulis*. This work has benefited from the support of the MMB Chair (Mathematics for Life and Environmental Sciences, “Chaire Modélisation Mathématique et Biodiversité”) VEOLIA-Ecole Polytechnique-MNHN-F.X.

## Statement of Authorship

All authors participated in the conception, analyses and interpretation of the model. All authors reviewed and edited the writing at all stages of revision.

## Appendices mathematical results

Our starting point is the replicator-mutator PDE

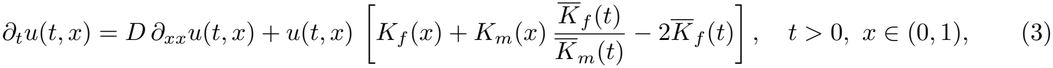

see subsection 2.2 for precise statements.

Our approach is mainly to analyze the stationary solutions to (3). This corresponds, for a given *D >* 0, to first formally letting *t* → +∞ in (3) to reach

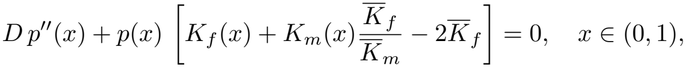

the equation for a stationary state *p* = *p*(*x*). In this framework, our mathematical results are stated in Appendix A and proved in Appendix B. After proving existence (and uniqueness) of a stationary solution in Proposition A.1, we focus on its number of local maxima (modes) as well as their localization. In particular we study the case *D* ≫ 1 in subsection A.3 and the case *D* → 0 in subsection A.4.

However, if we first formally let *D* → 0 in (3), we reach

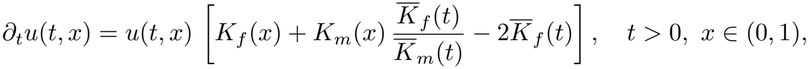

a model that we analyze in Appendix C.

Although mutation is absent from this equation, one may still introduce rare mutants as isolated events. Between successive mutation events, resident and mutant frequencies follow the resulting *D* = 0 dynamics. This separation underlies classical invasibility approaches, including the adaptive dynamics framework (Lesaffre et al., 2024), population genetics models (Charnov, 1979), and phenotypic models (Charnov et al., 1976). By contrast, our PDE describes mutation as a recurrent and continuous process.

### A Stationary solutions

#### A.1 Existence and uniqueness of the stationary state

Our first result concerns the existence and uniqueness of a stationary state for (1).

##### Proposition A.1

(Stationary state). *Assume that K*_*f*_, *K*_*m*_ ∈ *C*([0, 1]) *are nonnegative and not identically zero, with K*_*f*_ *nondecreasing and K*_*m*_ *nonincreasing. Then equation* (1) *admits a unique positive stationary state p satisfying*

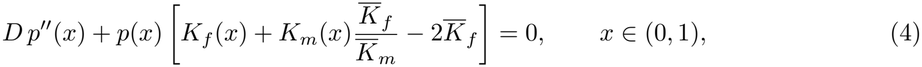

*with*

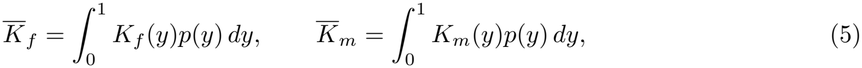

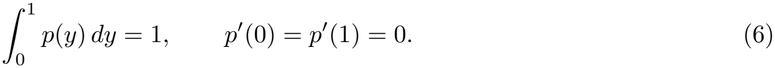

The next lemma shows that the ratio 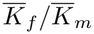 associated with the stationary state is bounded away from zero and infinity uniformly with respect to *D*.

##### Lemma A.2.

*Under the assumptions of Proposition A.1, let p be the unique positive stationary solution of* (1) *and set*

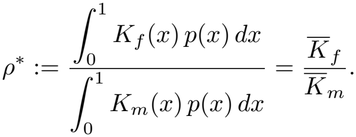

*Then ρ*^*\**^ *is bounded in* (0, +∞) *independently of D. More precisely:*

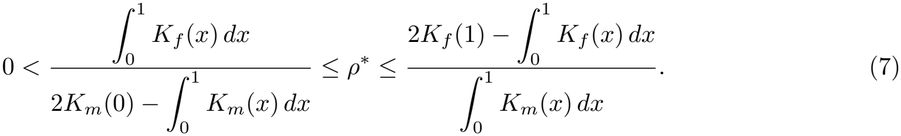

##### Remark A.3.

*In Proposition A.1, if we drop the monotonicity assumption of K*_*f*_ *and K*_*m*_, *we can still build a solution p, but uniqueness remains an open question. In the following, in order to remain as general as possible, Lemma A.4, Proposition A.5 and Proposition A.8 are stated for an arbitrary stationary state since they do not require the monotonicity assumption. By contrast, Proposition A.6, Conjecture A.7 and Theorem A.10 are stated for the unique stationary state of Proposition A.1 since they assume monotone fitness functions*.

In the rest of Appendix A our goal is to characterize the number of local maxima of the stationary state *p* as well as their localization in the range (0, 1). A first result tells us that there is at most one local maximum within (0, 1).

##### Lemma A.4.

*Let p be a stationary state for* (1) *solution of* (4)*–*(5)*–*(6). *Assume* 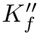 *and* 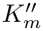 *are both positive (or negative) on* (0, 1). *Then p has at most one critical point in* (0, 1).

#### A.2 When fitness functions *K*_*f*_ and *K*_*m*_ are symmetric

In this section, we analyze the stationary distribution *p* without specifying the functional form of the fitness functions *K*_*f*_ and *K*_*m*_, but only supposing they are symmetric. We obtain general results about the number of modes of *p* and their localization for any mutational diffusivity *D*.

Assume

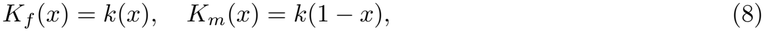

for some *k* : [0, 1] → [0, +∞). Denote (*p, λ*) the principal eigenpair of the Neumann operator *D∂*_*xx*_ + *K*_*f*_ (*x*) + *K*_*m*_(*x*), namely (*p, λ*) is the unique couple satisfying

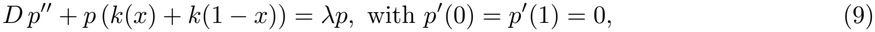

together with *p >* 0 in (0, 1), and the normalization condition 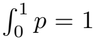. Since (*p*(1 − ·), *λ*) is “another” solution, we deduce that *p* = *p*(1 − ·), i.e. *p* is symmetric. Using this and integrating (9) over (0, 1), we infer that 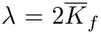 . Also 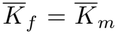. As a result the eigenfunction *p is* a (symmetric) stationary state for (1).

##### Proposition A.5

(Symmetric convex and symmetric concave cases). *Let the symmetry assumption* (8) *hold. Denote p the symmetric stationary state constructed above*.

1. *If k*^*′′*^ *<* 0 *on* (0, 1), *p is increasing on* (0, 1*/*2), *decreasing on* (1*/*2, 1), *i.e. unimodal with a unique local maximum at x* = 1*/*2.
2. *If k*^*′′*^ *>* 0 *on* (0, 1), *p is decreasing on* (0, 1*/*2), *increasing on* (1*/*2, 1), *i.e. bimodal with two local maxima at x* = 0 *and x* = 1.

In particular, Proposition A.5 encompasses *K*_*f*_ (*x*) = *x*^*γ*^ and *K*_*m*_(*x*) = (1 − *x*)^*γ*^, with 0 *< γ <* 1 or *γ >* 1, and is consistent with the numerical results in Fig. 3. This shows that *γ* = 1 is a threshold value separating two regimes: evolution leads either to a unimodal stationary distribution in (0, 1) with maximum at 1*/*2 or to a bimodal stationary distribution concentrated at the boundaries.

In the following we give results for the non-symmetric case for different mutation regimes (either *D* ≫ 1 or *D* → 0).

#### A.3 The strong mutation regime (*D* ≫ 1)

In this section, we study the stationary state in the regime *D* → +∞. We first show, for general fitness functions satisfying the assumptions of Proposition A.1, that the stationary density converges uniformly to the uniform density.

##### Proposition A.6

(Convergence to a uniform density as *D* → +∞). *Under the assumptions of Proposition A.1, let p*_*D*_ *be the unique normalized positive stationary state associated with D >* 0. *Then*

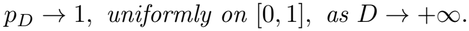

*More precisely*, 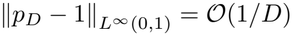 *as D* → +∞.

We then characterize the stationary distribution *p*_*D*_ for large but finite values of *D* when the fitness functions *K*_*f*_ and *K*_*m*_ are specified as power-law functions, yet not symmetric. The following is based on *formal* asymptotic expansions so we only state it as a conjecture sustained by numerical simulations in Fig. 3a and Fig. 5a.

##### Conjecture A.7

(*D* ≫ 1). *Assume*

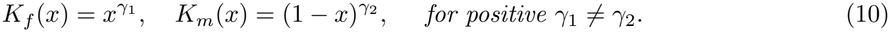

*Then, for D* ≫ 1, *the following holds*.

i. *If* 0 *< γ*_1_, *γ*_2_ *<* 1 *(concave-concave case), p is unimodal, reaching a unique local maximum at some x*_0_ ∈ (0, 1). *If γ*_1_ *> γ*_2_ *then* 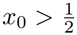 . *If γ*_1_ *< γ*_2_ *then* 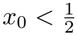.
ii. *If γ*_1_, *γ*_2_ *>* 1 *(convex-convex case), p is bimodal, reaching its two maxima at x* = 0 *and x* = 1.
iii. *If* 0 *< γ*_1_ *<* 1 *< γ*_2_ *(concave-convex case), p is unimodal reaching a unique local maximum at x* = 0.
iv. *If* 0 *< γ*_2_ *<* 1 *< γ*_1_ *(convex-concave case), p is unimodal reaching a unique local maximum at x* = 1.

These results show that the stationary state *p* can be classified in four categories depending on convexity of the functions *K*_*f*_ and *K*_*m*_, i.e. depending on the position of the parameters *γ*_1_ and *γ*_2_ relative to 1.

#### A.4 The weak mutational diffusion regime (*D* → 0)

In this section, we characterize the weak limits of stationary densities as the mutation coefficient *D* tends to zero. For each *ρ >* 0, we introduce the effective fitness function

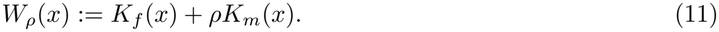

The parameter *ρ* will be determined self-consistently as the limiting ratio of mean female to mean male fitness. We first show, for general fitness functions *K*_*f*_ and *K*_*m*_, that any limiting measure is supported on the set of global maximizers of *W*_*ρ*_. In particular, when this set is finite, the limiting measure is a finite combination of Dirac masses. We then specialize to power-law fitness functions and explicitly identify the possible limiting measures, including the localizations and masses of their support points. We denote by *p*_*D*_ a stationary state associated with the mutation coefficient *D >* 0. We also write

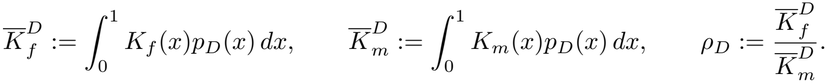

Thus *p*_*D*_ is the positive principal eigenfunction of the Neumann operator

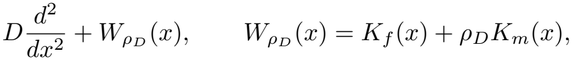

and the corresponding principal eigenvalue is

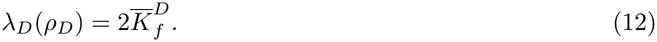

When *D* is small, the principal eigenfunction of 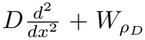 is expected to concentrate near the global maximizers of 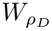 . The following proposition gives the precise necessary conditions satisfied by any weak limit of *p*_*D*_ as *D* → 0.

##### Proposition A.8

(Weak limit as *D* → 0). *Assume that, along a sequence D* → 0,

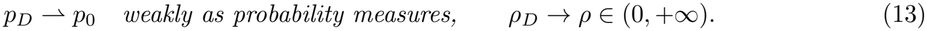

*Define*

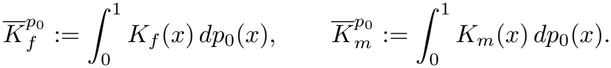

*Then*

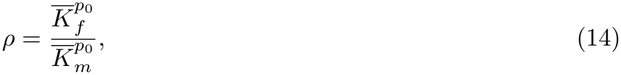

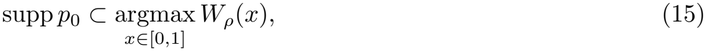

*and*

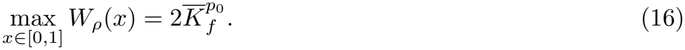

##### Remark A.9.

*The assumption that ρ*_*D*_ *converges to some ρ* ∈ (0, +∞) *is not restrictive for monotone fitness functions, see Lemma A.2*.

We now take advantage of Proposition A.8 to characterize the possible limiting measures when

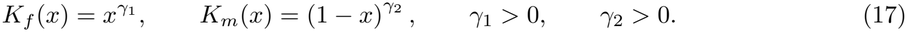

##### Theorem A.10

(Identifying the weak limits as *D* → 0). *Assume* (17), *with γ*_1_≠ 1 *and γ*_2_≠ 1. *Let p*_0_ *be a limiting measure satisfying the conclusions of Proposition A.8*. *Then the following alternatives fully describe p*_0_ *in terms of γ*_1_ *and γ*_2_.

i. *If* 0 *< γ*_1_, *γ*_2_ *<* 1 *(concave-concave case), then*

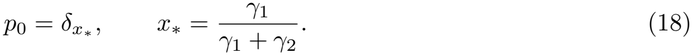
ii. *If γ*_1_, *γ*_2_ *>* 1 *(convex-convex case), then*

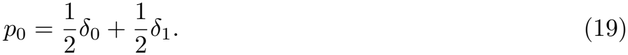
iii. *If* 0 *< γ*_1_ *<* 1 *< γ*_2_ *(concave-convex case), then there are two subcases. If*

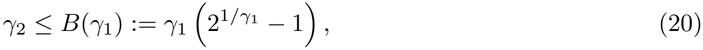

*then*

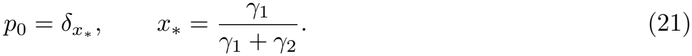

*If*

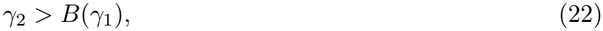

*then*

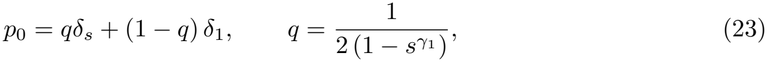

*where s* ∈ (0, 2^*−*1*/γ* 1^) *is the unique solution of*

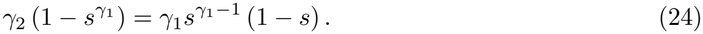

*At the transition γ*_2_ = *B*(*γ*_1_), *one has s* = 2^*−*1*/γ* 1^ *and q* = 1.
iv. *If* 0 *< γ*_2_ *<* 1 *< γ*_1_ *(convex-concave case), then there are two subcases. If*

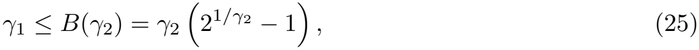

*then*

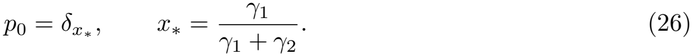

*If*

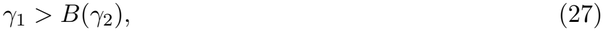

*then*

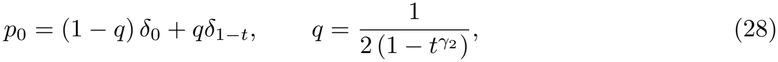

*where t* ∈ (0, 2^*−*1*/γ* 2^)*is the unique solution of*

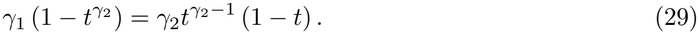

In the weak mutational diffusion regime (*D* → 0), the stationary distributions are either a single Dirac mass at 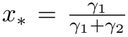 or a combination of two Dirac masses, one of which is located at a boundary. In summary, our results show that stationary distributions in the weak mutational diffusion regime can be split into four different categories.

i. Hermaphroditism: if 0 *< γ*_1_, *γ*_2_ *<* 1 (concave-concave case), and 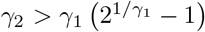 or if 0 *< γ*_1_ *<* 1 *< γ*_2_ (concave-convex case) and 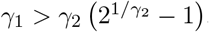. All individuals are hermaphroditic with trait value 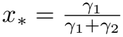 ;
ii. Dioecy: if *γ*_1_, *γ*_2_ *>* 1 (convex-convex case). The population is composed of individuals that are either pure females or pure males, with a balanced sex ratio 1 : 1.
iii. Gynodioecy: if 0 *< γ*_1_ *<* 1 *< γ*_2_ (concave-convex case) and 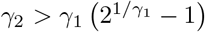. The population is composed of 1 − *q* pure females and *q* hermaphroditic individuals with trait *s* (see (23)–(24) for the values of *q* and *s*).
iv. Androdioecy: if 0 *< γ*_2_ *<* 1 *< γ*_1_ (convex-concave case) and 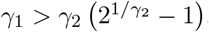. The population is composed of 1 − *q* pure males and *q* hermaphroditic individuals with trait 1 − *t* (see (28)–(29) for the values of *q* and *t*).

The above results are consistent with the numerical results in Fig. 3f and Fig. 5d.

#### A.5 Numerical computation of stationary states

The numerical method follows the spectral construction used in the proof of Proposition A.1. For fixed *D >* 0 and *ρ >* 0, we compute the normalized positive principal eigenpair (*φ*_*ρ*_, *λ*_*ρ*_) satisfying

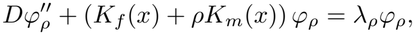

with

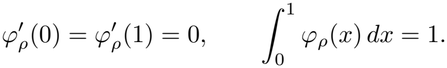

We then define

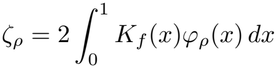

and determine the stationary value *ρ*_*D*_ by solving *λ*_*ρ*_ − *ζ*_*ρ*_ = 0. The stationary density is subsequently given by 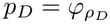 .

The interval [0, 1] is discretized into *N*_*x*_ = 400 uniform subintervals. The diffusion operator is approximated by second-order centered finite differences. For each value of *ρ*, the eigenvalues and eigenvectors of the resulting matrix are computed using `numpy.linalg.eig`. The principal eigenvalue is selected as the largest real eigenvalue, and the corresponding eigenvector is verified to have entries of positive sign and normalized using a discrete approximation of its integral.

The equation *λ*_*ρ*_ − *ζ*_*ρ*_ = 0 is solved with Brent’s method. Once *ρ*_*D*_ has been determined, the corresponding normalized principal eigenvector provides the numerical approximation of *p*_*D*_. This procedure directly targets the stationary problem and avoids approximating the stationary state by integrating (1) up to an arbitrarily chosen terminal time.

For the numerical phase diagrams, we consider

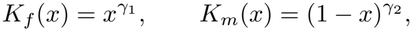

and sample 10^5^ parameter pairs (*γ*_1_, *γ*_2_) uniformly in [0, 4]^2^ for each considered value of *D*. Each stationary density is classified according to the number and location of its modes, including possible boundary modes. The corresponding principal eigenvalue is recorded as the stationary mean fitness.

The Jupyter notebooks implementing these computations and generating the manuscript figures are available in the Zenodo repository cited in the Data availability section.

### B Proofs of mathematical results of Appendix A

#### B.1 Existence and uniqueness of the stationary state

*Proof of Proposition A.1*. Let *λ* ∈ ℝ and *ρ >* 0. We look for positive solutions of

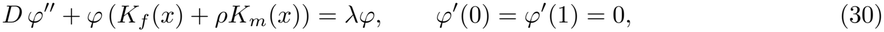

and

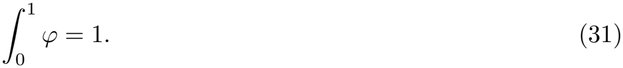

If (*λ, ρ, φ*) also fulfills the following relationships, then *φ* is a stationary solution of (1):

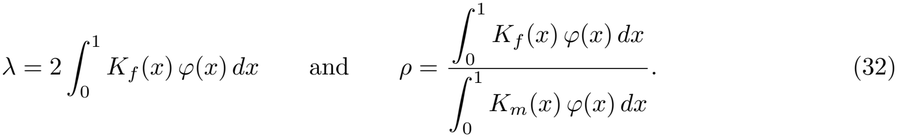

Krein-Rutman theory implies that, for each *ρ* ≥ 0, there is a unique couple (*φ*_*ρ*_, *λ*_*ρ*_) satisfying (30)-(31) and such that *φ*_*ρ*_ *>* 0 in (0, 1). The principal eigenvalue *λ*_*ρ*_ is characterized by the Rayleigh quotient:

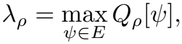

with

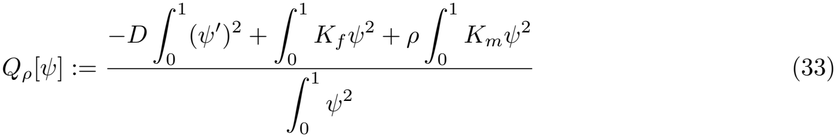

and

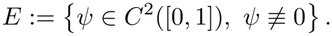

First, we observe that *ρ* ↦ *λ*_*ρ*_ is strictly increasing in [0, +∞). Indeed, if 0 ≤ *ρ*_1_ *< ρ*_2_, then

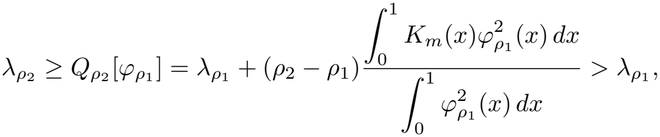

since *K* ≥ 0 is not identically zero and *φ >* 0. Also, since 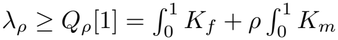, we have

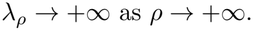

Next, for *ρ* = 0, integrating equation 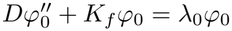 over (0, 1) and using the Neumann boundary conditions, we get

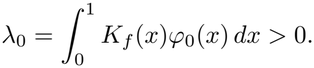

Now, set

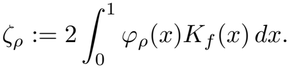

We claim that *ρ* ↦ *ζ*_*ρ*_ is nonincreasing on [0, +∞) (see below for the proof of this fact). Obviously *ζ*_0_ = 2*λ*_0_ and, since *K*_*f*_ is nondecreasing, *ζ*_*ρ*_ ≤ 2*K*_*f*_ (1) for all *ρ* ≥ 0.

From the above, ℱ(*ρ*) : *ρ* ↦ *λ*_*ρ*_ − *ζ*_*ρ*_ is continuous (classically, the continuity of *ρ* ↦ *λ*_*ρ*_ follows from the variational characterization (33), while that of *ρ* ↦ *φ*_*ρ*_ w.r.t. the *L*^*∞*^ norm follows from the fact that the principal eigenvalue is simple and normalized, and hence *ρ* ↦ *ζ*_*ρ*_ is continuous) and strictly increasing on [0, +∞), ℱ(0) = −*λ*_0_ *<* 0 and lim_*ρ→*+*∞*_ ℱ(*ρ*) = +∞. As a result there exists a unique *ρ*^*\**^ *>* 0 such that ℱ(*ρ*^*\**^) = 0, that is,

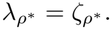

Thus, 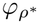 satisfies the first equality in (32). Integrating (30), we obtain

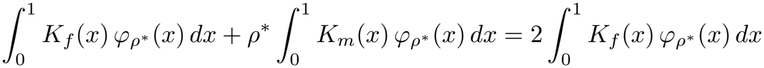

which in turn implies that the second equality in (32) is also satisfied. Thus, 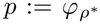 is a stationary solution of (1).

To prove uniqueness, let *φ* be any positive stationary solution satisfying 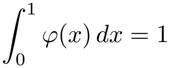, and set

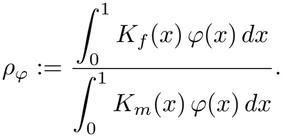

The stationary solution *φ* is a positive eigenfunction of (30) with *ρ* = *ρ*_*φ*_ and eigenvalue 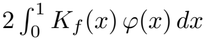. Hence, by uniqueness of the normalized principal eigenfunction,

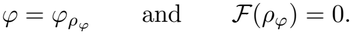

Since *ρ*^*\**^ is the unique zero of ℱ, it follows that

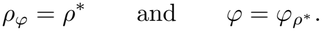

This proves the uniqueness of the stationary solution.□

*Proof of: ρ* ↦ *ζ*_*ρ*_ *is nonincreasing on* [0, +∞). Fix 0 ≤ *ρ*_1_ *< ρ*_2_ and set

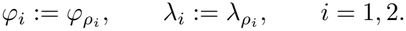

Define

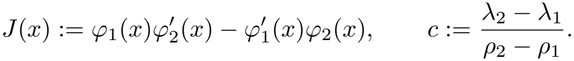

Multiplying the equation for *φ*_2_ by *φ*_1_, the equation for *φ*_1_ by *φ*_2_, and subtracting, we obtain

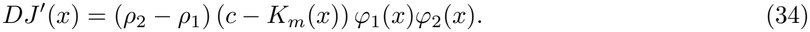

The Neumann boundary conditions imply *J*(0) = *J*(1) = 0 so that, integrating (34) over (0, 1),

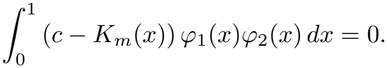

The function *c* − *K*_*m*_ being nondecreasing, this implies that there exists *x*_*m*_ ∈ (0, 1) such that

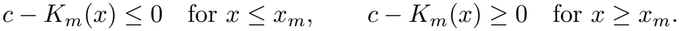

Therefore, *J*^*′*^ is first nonpositive and then nonnegative. Since *J*(0) = *J*(1) = 0, we deduce that

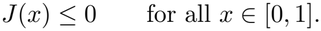

Set

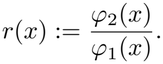

Since 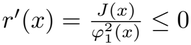, *r* is nonincreasing. Moreover, the normalization of the eigenfunctions gives

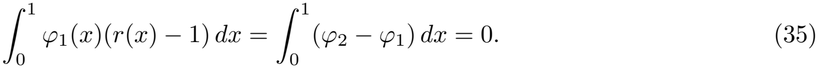

Hence, there exists *x*_0_ ∈ (0, 1) such that

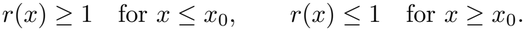

Using (35) we obtain

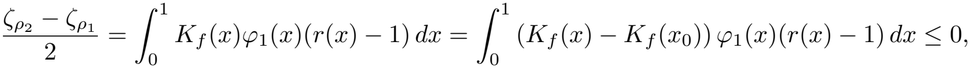

since the two factors

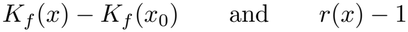

have opposite signs on both sides of *x*_0_ because *K*_*f*_ is nondecreasing. Thus, *ζ*_*ρ* 2_ ≤ *ζ*_*ρ* 1_ .□

*Proof of Lemma A.2*. Using the notations in the proof of Proposition A.1, the unique stationary solution is 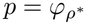 and

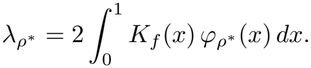

Testing the Rayleigh quotient (33) with *ρ* = *ρ*^*\**^ and *ψ* ≡ 1 gives

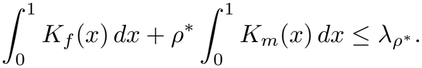

Since 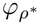 is positive, normalized by 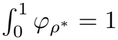, and *K*_*f*_ is nondecreasing, we have

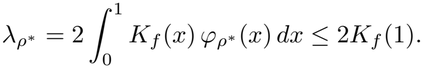

Consequently,

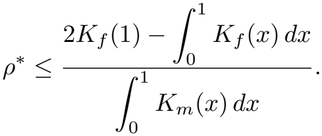

To obtain the lower bound, divide (30) by *ρ*^*\**^. Then 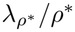 is the principal eigenvalue associated with the operator (with Neumann boundary conditions)

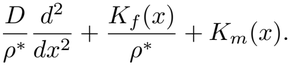

Testing the corresponding Rayleigh quotient with the constant function *ψ* ≡ 1 gives

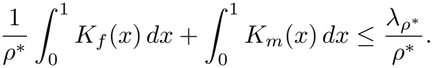

Moreover, integrating (30) and using 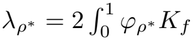, we obtain

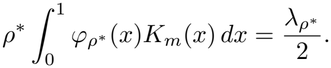

Therefore,

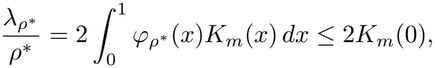

because *K*_*m*_ is nonincreasing. It follows that

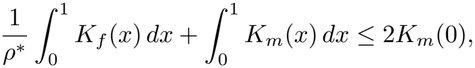

and hence

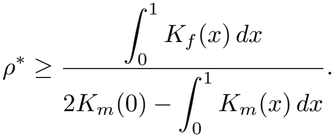

This proves (7).□

*Proof of Lemma A.4*. We have

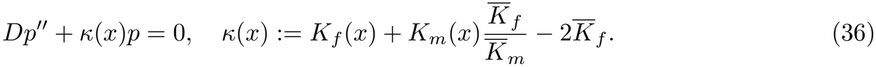

Assume *p* has *n* interior critical points, namely *x*_0_ := 0 *< x*_1_ *<* · · · *x*_*n*_ *<* 1 := *x*_*n*+1_. Integrating (36) over (*x*_*i*_, *x*_*i*+1_), we obtain

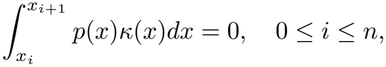

so that *κ* has to change sign at least *n* + 1 times. Since *κ* is strictly convex (or concave) on (0, 1) (see assumptions of the lemma), *n* + 1 ≤ 2.

#### B.2 The symmetric case

*Proof of Proposition A.5*. We have

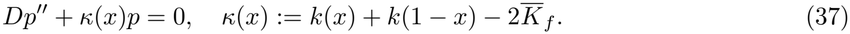

Integrating (37), we get 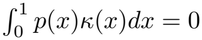 so *κ* must change sign at least once on (0, 1).

Assume *k*^*′′*^ *<* 0 or *k*^*′′*^ *>* 0 on (0, 1). It follows from Lemma A.4 and the symmetry of *p* that *p* has exactly one interior critical point, at 1*/*2.

Now, assume *k*^*′′*^ *<* 0 on (0, 1). Function *κ* is symmetric, strictly concave, and must change sign at least once on (0, 1). As a result, for some *x*^*\**^ ∈ (0, 1*/*2), *κ <* 0 on (0, *x*^*\**^) ∪ (1 − *x*^*\**^, 1) while *κ >* 0 on (*x*^*\**^, 1 − *x*^*\**^). It follows from (37) that *p* is strictly convex on (0, *x*^*\**^) ∪ (1 − *x*^*\**^, 1) and strictly concave on (*x*^*\**^, 1 − *x*^*\**^). Putting all together, we have *p*^*′*^ *>* 0 on (0, 1*/*2), *p*^*′*^ *<* 0 on (1*/*2, 1), which proves (*i*). The proof of (*ii*) is similar and omitted.

#### B.3 The regime *D* ≫ 1

*Proof of Proposition A.6*. As in the proof of Lemma A.4, set

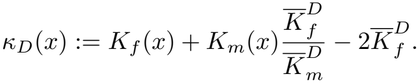

Lemma A.2 and the normalization 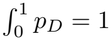 imply that

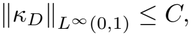

for some constant *C* independent of *D*. Integrating the equation 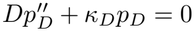 between 0 and *x* and using 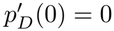, we get

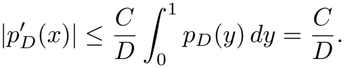

Finally, using again the normalization 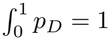, we obtain, for every *x* ∈ [0, 1],

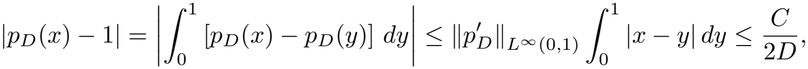

which concludes the proof.

*Formal asymptotic expansions sustaining Conjecture A.7*. We denote *ε* := *D*^*−*1^ so that

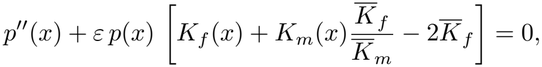

where *p* = *p*_*ε*_. As *ε* → 0, we perform formal asymptotic expansions and write

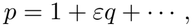

so that

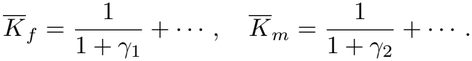

Plugging these expansions into the equation, retaining the terms of order *ε*, we reach

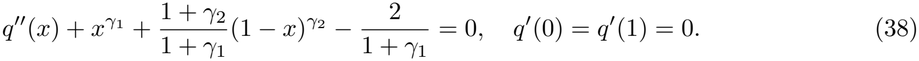

In particular 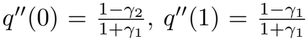 so that there is a maximum at 0, resp. 1, as soon as *γ*_2_ *>* 1, resp. *γ*_1_ *>* 1 (and a minimum when *>* is replaced by *<*).

i. In the concave-concave case (0 *< γ*_1_, *γ*_2_ *<* 1), it therefore follows from the above and Lemma A.4 that *p* is unimodal, reaching a unique maximum at some *x*_0_ ∈ (0, 1). Where is this optimum *x*_0_? Integrating (38), we get
ii. 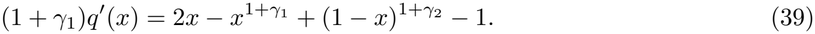

Obviously when *γ*_1_ = *γ*_2_, 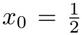 . Now, if we let *γ*_2_ = *γ, γ*_1_ = *γ* + *h* and apply the implicit function theorem to *f* (*h, x*) = 2*x* − *x*^1+*γ*+*h*^ + (1 − *x*)^1+*γ*^ − 1, we obtain

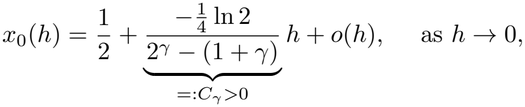

meaning that, slightly “moving” downward the diagonal *γ*_1_ = *γ*_2_, the point is moving towards 1. Note also that *C*_*γ*_ → +∞ as *γ* → 0 or *γ* → 1.
iii. In the convex-convex case (*γ*_1_, *γ*_2_ *>* 1), it therefore follows from the above and Lemma A.4 that *p* is bimodal, reaching its two maxima at *x* = 0 and *x* = 1.
iv. In the concave-convex case (0 *< γ*_1_ *<* 1 *< γ*_2_), it therefore follows from the above and Lemma A.4 that *p* is unimodal, reaching its unique maximum at *x* = 0.
v. The convex-concave case is studied similarly.

#### B.4 The regime *D* → 0

*Proof of Proposition A.8*. For fixed *ρ >* 0, the Neumann principal eigenvalue of 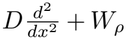 is characterized by the Rayleigh formula

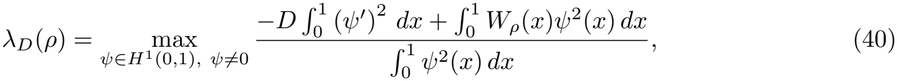

from which we collect the upper bound

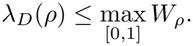

Conversely, let *x*_0_ be a global maximizer of *W*_*ρ*_. For every *η >* 0, we may choose a nonzero test function *ψ*_*η*_ ∈ *H*^1^(0, 1), concentrated in a sufficiently small neighborhood of *x*_0_, such that

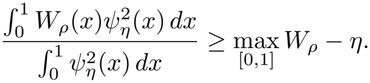

Using *ψ*_*η*_ in the Rayleigh quotient gives

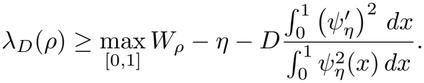

For fixed *η*, we let *D* → 0, and then let *η* → 0. Therefore

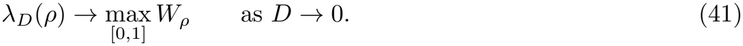

Now assume that *ρ*_*D*_ → *ρ*. Since *W*_*ρ D*_ → *W*_*ρ*_ uniformly on [0, 1], we obtain

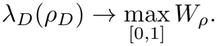

Using (12), this yields

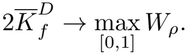

By weak convergence of *p*_*D*_ and continuity of *K*_*f*_ and *K*_*m*_, we also have

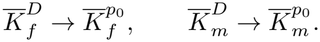

Since 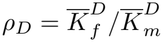 and *ρ*_*D*_ → *ρ* ∈ (0, +∞), we obtain

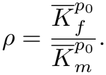

Moreover,

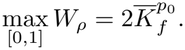

This proves (14) and (16).

It remains to identify the support of *p*_0_. Using (14), we compute

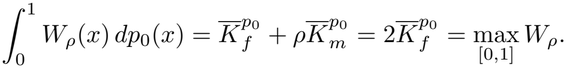

Since *W*_*ρ*_(*x*) ≤ max *W*_*ρ*_ for all *x* ∈ [0, 1], equality of the average with the maximum implies that *W*_*ρ*_ = max *W*_*ρ*_ *p*_0_-almost everywhere. Indeed, if *M* := max *W*_*ρ*_, then *M* − *W*_*ρ*_ ≥ 0 and

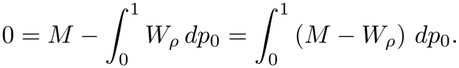

Hence *M* − *W*_*ρ*_ = 0 *p*_0_-almost everywhere. Since *W*_*ρ*_ is continuous, the maximizer set is closed, and therefore

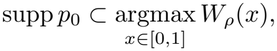

which concludes the proof.

*Proof of Theorem A.10*. The proof mainly relies on the three conclusions of Proposition A.8, namely

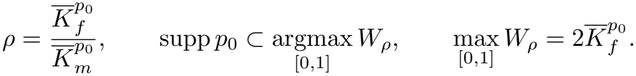

Since *W*_*p*_(1)=1, 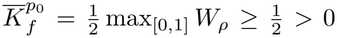. As 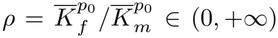 we also have 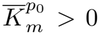. Hence *p*_0_ cannot be supported only at 0 or only at 1.

##### Case (*i*)

0 *< γ*_1_, *γ*_2_ *<* 1. Since *ρ >* 0, the function *W*_*ρ*_(*x*) = *x*^*γ* 1^ + *ρ* (1 − *x*)^*γ* 2^ is strictly concave on (0, 1). Moreover,

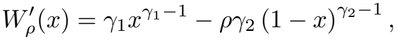

so 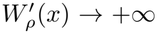as *x* → 0 and 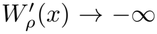 as *x* → 1. Therefore *W*_*ρ*_ has a unique global maximizer, denoted by *x*_*\**_ ∈ (0, 1). Since supp *p*_0_ is contained in the maximizer set of *W*_*ρ*_, we must have

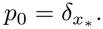

Then

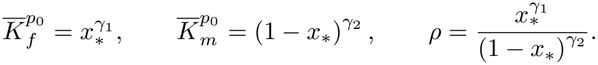

Since *x*_*\**_ is an interior maximizer, 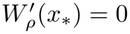, namely

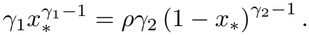

Substituting the expression for *ρ* gives

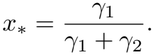

This proves (18).

##### Case (*ii*)

*γ*_1_, *γ*_2_ *>* 1. Since *ρ >* 0, the function *W*_*ρ*_(*x*) = *x*^*γ* 1^ + *ρ* (1 − *x*)^*γ* 2^ is strictly convex on (0, 1). Hence its maximum over [0, 1] is reached at the endpoints. Thus

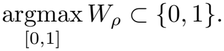

Since the measure *p*_0_ cannot be supported only at 0 or only at 1, it follows from supp *p*_0_ ⊂ argmax[0,1] *W*_*ρ*_ that both endpoints must be maximizers:

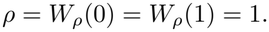

We may therefore write

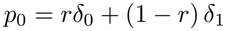

with *r* ∈ (0, 1). Then

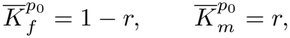

and the identity 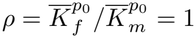 enforces *r* = 1*/*2. As a result

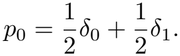

This proves (19).

##### Case (*iii*)

0 *< γ*_1_ *<* 1 *< γ*_2_. We have

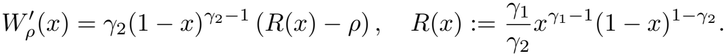

The sign of 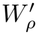 on (0, 1) is that of *R*(*x*) − *ρ*. We observe that *R* decreases on (0, *x*_*R*_) and increases on (*x*_*R*_, 1), with

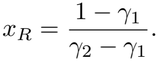

Moreover,

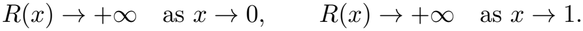

Therefore the equation *R*(*x*) = *ρ* has either no solution, one double solution, or two solutions. Hence *W*_*ρ*_ has at most two interior critical points, and thus at most one interior local maximum.

Moreover, *x* = 0 cannot be a global maximizer, because *W*_*ρ*_ is increasing immediately to the right of The only endpoint that can be a global maximizer is therefore *x* = 1. Hence the global maximizer set of *W*_*ρ*_ is either a single interior point, the endpoint 1, or one interior point together with the endpoint 1. We recall that the support cannot be reduced to {1}. Therefore only two possibilities remain.

- *First possibility: one interior support point*. Assume first that

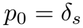

for some *s* ∈ (0, 1). As in Case (*i*), the condition 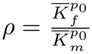 gives

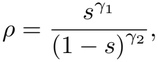

and the interior critical point condition 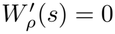 gives

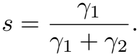

For this point to be a global maximizer, it is necessary that its height is at least the height of the endpoint 1. Since *W*_*ρ*_(*s*) = 2*s*^*γ* 1^, *W*_*ρ*_(1) = 1, we need 2*s*^*γ* 1^ ≥ 1, that is

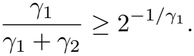

Solving for *γ*_2_ gives

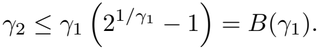

This is precisely the condition (20). Therefore, if the limit is a one-point measure in the concave-convex regime, it must be

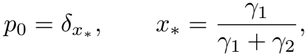

and this can occur only when *γ*_2_ ≤ *B*(*γ*_1_).
- *Second possibility: one interior support point and the endpoint* 1. Assume now that

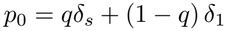

with *s* ∈ (0, 1) and *q* ∈ (0, 1). Since both *s* and 1 belong to the maximizer set of *W*_*ρ*_, their heights are equal:

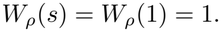

Therefore *s*^*γ* 1^ + *ρ* (1 − *s*)^*γ* 2^ = 1, or equivalently

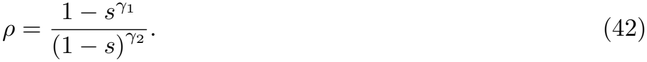

Moreover, since *s* is an interior maximizer, 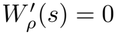, namely

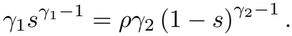

Substituting (42), we obtain 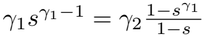, which is equivalent to

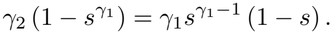

This is (24).

We now determine the mass *q*. Since *W*_*ρ*_(1) = 1, one has 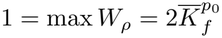 . Thus

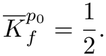

On the other hand,

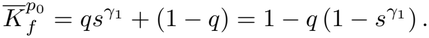

As a result, 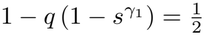, and hence

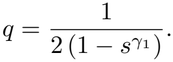

For this to define a probability mass with *q* ≤ 1, we need 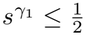, or equivalently

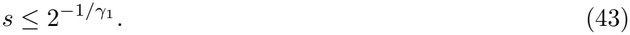

It remains to show that, when *γ*_2_ *> B*(*γ*_1_), equation (24) has exactly one solution satisfying (43).

Define

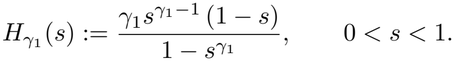

Then (24) is equivalent to

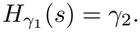

First, we note that *H*_*γ* 1_ is strictly decreasing on (0, 1). Moreover,

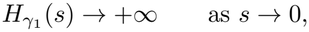

and, for *s*_0_ := 2^*−*1*/γ* 1^, we have

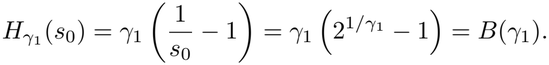

Therefore, if *γ*_2_ *> B*(*γ*_1_), there exists a unique solution

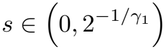

of (24). This gives the measure

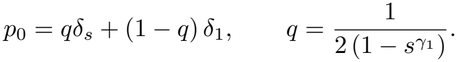

If *γ*_2_ = *B*(*γ*_1_), then the unique solution is 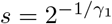, and the above formula gives *q* = 1.

##### Case (*iv*)

0 *< γ*_2_ *<* 1 *< γ*_1_. This case is symmetric to the previous one under the change of variable *y* = 1 − *x* and the exchange of *γ*_1_ and *γ*_2_.

### C No mutation regime (*D* = 0)

In this section, as explained above, we first let *D* → 0 in (3) and consider

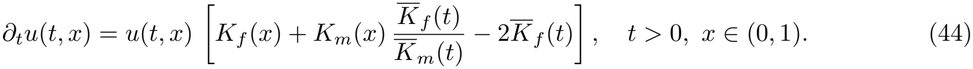

In the absence of mutation, *any* Dirac mass at *any x*_0_ ∈ (0, 1) is a stationary solution to (44). In order to specify the evolutionarily relevant ones, we consider the dynamics of a resident trait *x*_0_ ∈ (0, 1) and a mutant trait *x*_1_ ∈ [0, 1] \ {*x*_0_} given as

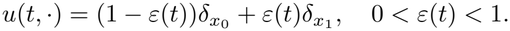

Plugging the change in the distribution

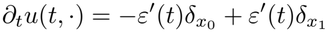

into (44), and identifying the coefficients of 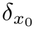 and 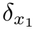, we obtain

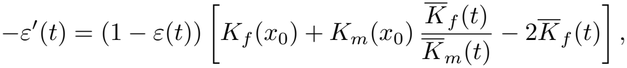

and

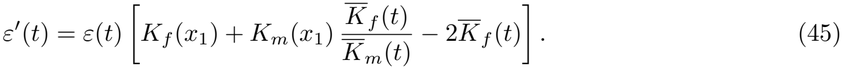

Summing these two identities and using the second one, we obtain

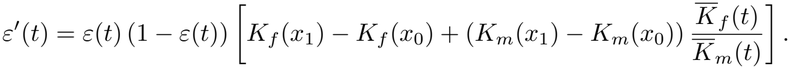

Using

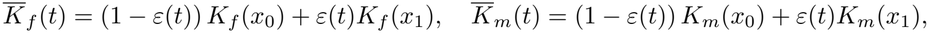

we therefore obtain

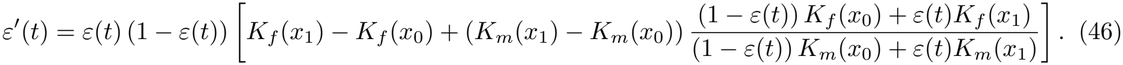

#### C.1 Rare mutant close to the resident: recovering the criterion in Lesaffre et al. (2024)

We here assume that the mutant trait is initially rare, that is

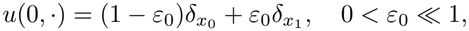

and close to the resident trait, that is

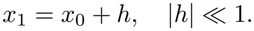

At least for short times, the mutant trait remains rare so that 0 *< ε*(*t*) ≪ 1. Hence, we may write

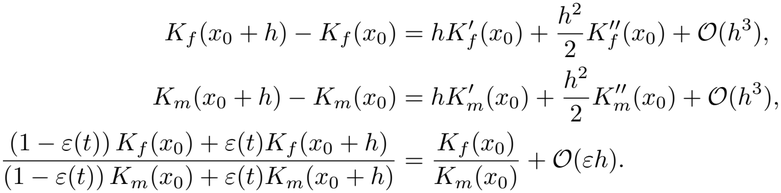

Plugging these expansions into (46) yields

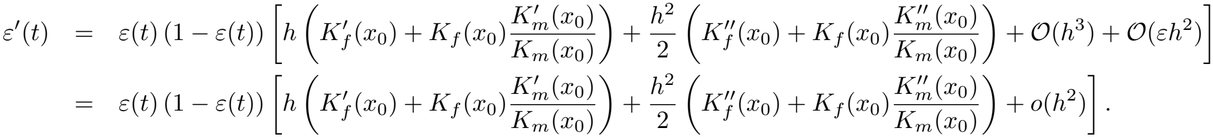

Hence, the Dirac mass at *x*_0_ = *x*^*\**^ is locally evolutionarily stable if

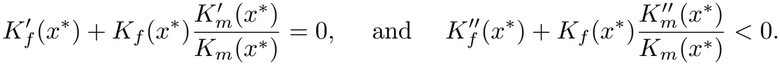

When the fitness functions are specified as power-law functions as in (10), these conditions reduce to

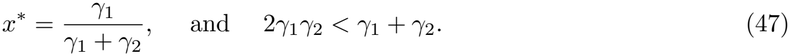

We thus recover the results of (Lesaffre et al., 2024, Section B.1.2.1) who derived these conditions using the adaptive dynamics framework.

#### C.2 Stability of *x*^*\**^ against a rare mutant (not necessarily close): recovering the criterion in Charnov (1979)

Still considering power-law fitness functions (10), we consider a resident trait at 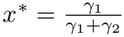 and investigate whether it can be invaded by *any* rare mutant trait *x*_1_ (not necessarily close to *x*^*\**^). To do so, we return to (45), which provides

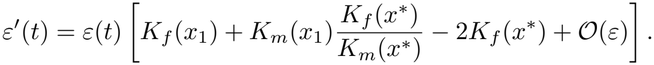

Hence, the condition for *x*^*\**^ to be invasible by a rare mutant *x*_1_ is

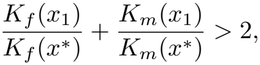

which is the condition obtained by Charnov (1979). Defining

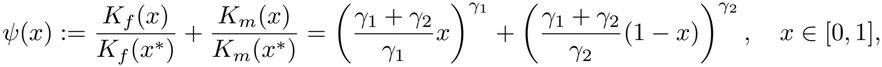

we are left to finding positive parameters *γ*_1_, *γ*_2_ so that max_0*≤x≤*1_ *ψ*(*x*) *>* 2. Below we disregard the singular case *γ*_1_ = *γ*_2_ = 1.

- If 0 *< γ*_1_, *γ*_2_ ≤ 1, then *ψ* is strictly concave on (0, 1). Moreover, *ψ*^*′*^(*x*^*\**^) = 0 and *ψ*(*x*^*\**^) = 2. Hence, *x*^*\**^ is a global maximizer of *ψ* and max_*x∈*[0,1]_ *ψ*(*x*) = *ψ*(*x*^*\**^) = 2. Therefore, no rare mutant can invade *x*^*\**^.
- If *γ*_1_, *γ*_2_ ≥ 1, then *ψ* is strictly convex on (0, 1) so that max_*x∈*[0,1]_ *ψ*(*x*) = max{*ψ*(0), *ψ*(1)}. If *γ*_1_ *>* 1, then

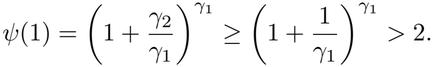

Similarly, if *γ*_2_ *>* 1, then *ψ*(0) *>* 2. Thus, there exists a rare mutant that can invade *x*^*\**^.

- If 0 *< γ*_1_ *<* 1 *< γ*_2_, then *ψ*^*′′*^ may change sign. However, *ψ*^*′*^(*x*) = 0 is equivalent to

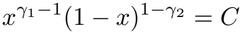

for some *C* = *C*(*γ*_1_, *γ*_2_) *>* 0. The function on the left-hand side tends to +∞ as *x* → 0 and as *x* → 1, and has a unique minimum. Hence, *ψ*^*′*^ has at most two zeros (one being *x*^*\**^). Since *ψ*^*′*^(0) = +∞ and *ψ*^*′*^(1) *>* 0, these properties imply

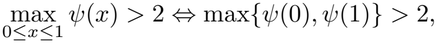

as one can easily check. As result

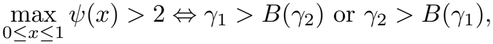

where *B*(*x*) := *x*(2^1*/x*^ − 1) for *x >* 0. Since *B* : (0, +∞) → (ln 2, +∞) is a decreasing bijection we are left with

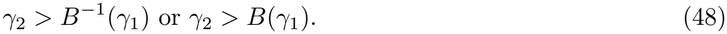

We claim that

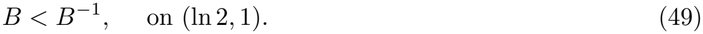

Indeed define *g*(*x*) := *B*(*B*(*x*)) − *x* for *x* ∈ (ln 2, 1). Observe that *g*(1) = 0. Since −1 *< B*^*′*^(*u*) *<* 0 for any *u >* ln 2, we have *g*^*′*^(*x*) = *B*^*′*^(*B*(*x*))*B*^*′*^(*x*) − 1 *<* 0 so that *g*(*x*) *>* 0 for *x* ∈ (ln 2, 1), which proves (49). Hence, in the considered range 0 *< γ*_1_ *<* 1 *< γ*_2_, condition (48), in view of (49), reduces to

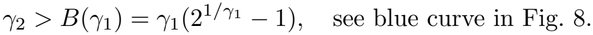

- If 0 *< γ*_2_ *<* 1 *< γ*_1_ then, by symmetry, we find

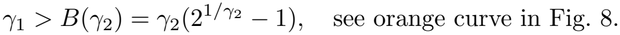

##### Remark C.1.

*In the* (*γ*_1_, *γ*_2_) *parameter plane, see Fig. 8, the bimodal region corresponding to gynodioecy and androdioecy is larger under Charnov’s criterion because invasion is considered from any point in the phenotypic space (Charnov, 1979), whereas Lesaffre et al.’s criterion only considers invasions by nearby mutants (Lesaffre et al., 2024)*. *This difference reflects the distinction between global and local invasibility*.

**Figure 8:**
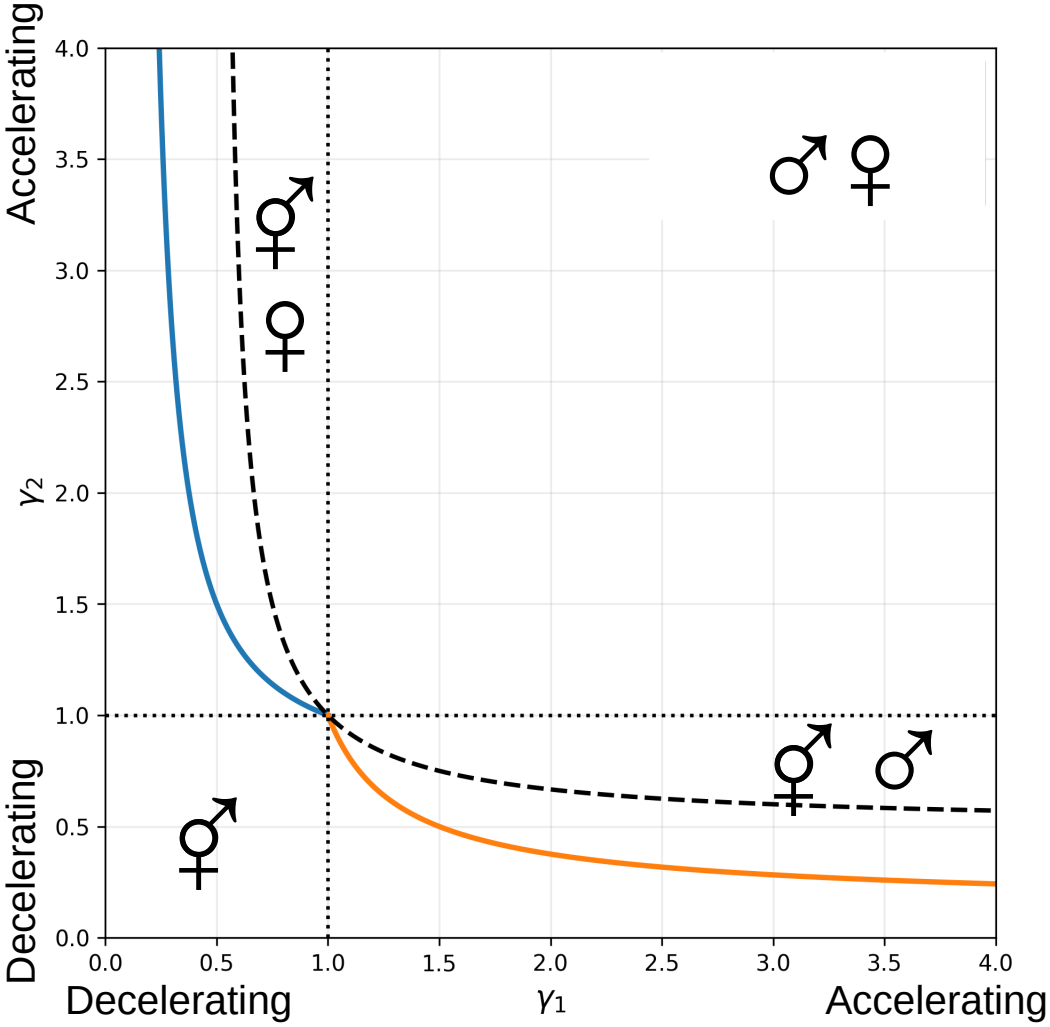
evolutionarily stable sex allocation predicted by invasion of a rare mutant, for power-law fitness functions 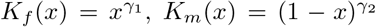. Dioecy: square delimited by *γ*_1_ ≥ 1 and *γ*_2_ ≥ 1 (dotted lines). Hermaphroditism: region below the dashed hyperbola 2*γ*_1_*γ*_2_ = *γ*_1_ + *γ*_2_ if only nearby mutants are considered (see subsection C.1, Lesaffre et al. 2024), or region below the thick blue curve *γ*_2_ = *γ*_1_ 2^1*/γ*1^ − 1 and the thick orange curve *γ*_1_ = *γ*_2_ 2^1*/γ*2^ − 1 if all mutants are considered (see subsection C.2, Charnov 1979). Gynodioecy and androdioecy are predicted between the hermaphroditism and the dioecy regions.

## References

Sally Adams, Sophie Tandonnet, and Andre Pires-daSilva. Balancing selfing and outcrossing: the genetics and cell biology of nematodes with three sexual morphs. Genetics, 229(2):iyae173, 2025.

Matthieu Alfaro and Rémi Carles. Explicit solutions for replicator-mutator equations: extinction versus acceleration. SIAM J. Appl. Math., 74(6):1919–1934, 2014. ISSN 0036-1399. doi: 10.1137/140979411. URL https://doi.org/10.1137/140979411.

Matthieu Alfaro and Rémi Carles. Replicator-mutator equations with quadratic fitness. Proc. Amer. Math. Soc., 145(12):5315–5327, 2017. ISSN 0002-9939. doi: 10.1090/proc/13669. URL https://doi.org/10.1090/proc/13669.

Matthieu Alfaro and Mario Veruete. Evolutionary branching via replicator-mutator equations. Journal of Dynamics and Differential Equations, 31(4):2029–2052, 2019. doi: 10.1007/s10884-018-9692-9.

Matthieu Alfaro, François Hamel, Florian Patout, and Lionel Roques. Adaptation in a heterogeneous environment. II: To be three or not to be. Journal of Mathematical Biology, 87(5):68, 2023. doi: 10.1007/s00285-023-01996-4.

Matthieu Alfaro, Florian Lavigne, and Lionel Roques. A host-pathogen coevolution model, part I: Run straight for your life. Nonlinear Analysis: Real World Applications, 84:104305, 2025. doi: 10.1016/j.nonrwa.2024.104305.

Tia-Lynn Ashman. Constraints on the evolution of males and sexual dimorphism: field estimates of genetic architecture of reproductive traits in three populations of gynodioecious fragaria virginiana. Evolution, 57(9):2012–2025, 2003.

TL Ashman and CJ Majetic. Genetic constraints on floral evolution: a review and evaluation of patterns. Heredity, 96(5):343–352, 2006.

Estelle Barbot, Mathilde Dufaÿ, Jeanne Tonnabel, Cécile Godé, and Isabelle De Cauwer. On the function of flower number: disentangling fertility from pollinator-mediated selection. Proceedings of the Royal Society B: Biological Sciences, 289(1987):20221987, 2022.

Nicholas H. Barton, Alison M. Etheridge, and Amandine Véber. The infinitesimal model: Definition, derivation, and implications. Theoretical Population Biology, 118:50–73, 2017. doi: 10.1016/j.tpb.2017.06.001.

Steven B Broyles and Robert Wyatt. Paternity analysis in a natural population of asclepias exaltata: multiple paternity, functional gender, and the “pollen-donation hypothesis”. Evolution, 44(6):1454–1468, 1990.

Reinhard Bürger. On the maintenance of genetic variation: global analysis of Kimura’s continuum-of-alleles model. J. Math. Biol., 24:341–351, 1986.

N Champagnat, R Ferriére, and S Méléard. Unifying evolutionary dynamics: from individual stochastic processes to macroscopic models. Theoretical Population Biology, 69(3):297–321, 2006.

Nicolas Champagnat, Régis Ferriére, and Sylvie Méléard. From individual stochastic processes to macro-scopic models in adaptive evolution. Stoch. Models, 24:2–44, 2008. ISSN 1532-6349,1532-4214. doi: 10.1080/15326340802437710. URL https://doi.org/10.1080/15326340802437710.

Eric L Charnov. The genetical evolution of patterns of sexuality: Darwinian fitness. The American Naturalist, 113(4):465–480, 1979.

Eric L Charnov. The theory of sex allocation, volume 18. Princeton university press, 1982.

Eric L Charnov, James J Bull, and J Maynard Smith. Why be an hermaphrodite? Nature, 263(5573): 125–126, 1976.

Kai-Hsiu Chen and John R Pannell. Mapping fitness landscapes to interpret sex allocation in hermaphrodites. Current Biology, 35(10):2354–2364, 2025.

Pierre Collet, Sylvie Méléard, and Johan A. J. Metz. A rigorous model study of the adaptive dynamics of mendelian diploids. Journal of Mathematical Biology, 67(3):569–607, 2013. doi: 10.1007/s00285-012-0562-5.

Guillaume G Cossard and John R Pannell. A functional decomposition of sex inconstancy in the dioecious, colonizing plant mercurialis annua. American Journal of Botany, 106(5):722–732, 2019.

Léonard Dekens and Sepideh Mirrahimi. Dynamics of dirac concentrations in the evolution of quantitative alleles with sexual reproduction. Nonlinearity, 35(11):5781–5812, 2022. doi: 10.1088/1361-6544/ac91bb.

Lynda F Delph and David G Lloyd. Environmental and genetic control of gender in the dimorphic shrub hebe subalpina. Evolution, pages 1957–1964, 1991.

Wagner Luiz dos Santos, Kátia Cavalcanti Pôrto, and Fábio Pinheiro. An overview of reproductiveallocation and reproductive costs in bryophytes: Challenges and prospects: Wl dos santos et al. The Botanical Review, 90(2):130–159, 2024.

Wagner Luiz Dos Santos, Adaíses Simone Maciel-Silva, Fábio Pinheiro, and Rose Marks. The architecture of sexual systems in mosses and its ecological and evolutionary consequences. Integrative and Comparative Biology, 66:icag016, 2026.

Bodil K Ehlers and Thomas Bataillon. ‘inconstant males’ and the maintenance of labile sex expression in subdioecious plants. New Phytologist, 174(1):194–211, 2007.

Ronald Aylmer Fisher. The Genetical Theory of Natural Selection. The Clarendon Press, Oxford, UK, 1930.

Theodore H Fleming, Sandrine Maurice, and JL Hamrick. Geographic variation in the breeding system and the evolutionary stability of trioecy in pachycereus pringlei (cactaceae). Evolutionary Ecology, 12 (3):279–289, 1998.

Alexandre Francq, Pierre Saumitou-Laprade, Philippe Vernet, and Sylvain Billiard. A paradigm shift, or a paradigm adjustment? the evolution of the oleaceae mating system as a small-scale kuhnian case study. The Quarterly Review of Biology, 98(2):61–83, 2023.

Jörn F Gerchen, Nora Villamil-Buenrostro, Xinji Li, Ehouarn Le Faou, and John R Pannell. Sex-allocation trade-offs and their genetic architecture revealed by experimental evolution. Proceedings of the National Academy of Sciences, 123(2):e2427240123, 2026.

Marie-Eve Gil, François Hamel, Guillaume Martin, and Lionel Roques. Dynamics of fitness distributions in the presence of a phenotypic optimum: an integro-differential approach. Nonlinearity, 32(10):3485–3522, 2019. doi: 10.1088/1361-6544/ab1bbe.

VN Godin. Trioecy in flowering plants. In Doklady Biological Sciences, volume 507, pages 301–311. Springer, 2022.

Francois Hamel, Florian Lavigne, Guillaume Martin, and Lionel Roques. Dynamics of adaptation in an anisotropic phenotype-fitness landscape. Nonlinear Analysis: Real World Applications, 54:103107, 2020. doi: 10.1016/j.nonrwa.2020.103107.

Francois Hamel, Florian Lavigne, and Lionel Roques. Adaptation in a heterogeneous environment. I: Persistence versus extinction. Journal of Mathematical Biology, 83(2):14, 2021. doi: 10.1007/s00285-021-01637-8.

Camille Jolivel, Nina Joffard, Cécile Godé, Eric Schmitt, and Isabelle De Cauwer. Male-biased sexual selection persists across contrasting habitats in a dioecious plant. bioRxiv, pages 2026–02, 2026.

Jos Käfer, Marcos Méndez, and Sylvain Mousset. Labile sex expression in angiosperm species with sex chromosomes. Philosophical Transactions of the Royal Society B: Biological Sciences, 377(1850): 20210216, 2022.

Motoo Kimura. A stochastic model concerning the maintenance of genetic variability in quantitative characters. Proc. Natl. Acad. Sciences, 54(3):731–736, 1965.

Mark Kirkpatrick and Nicholas H Barton. Evolution of a species’ range. The American Naturalist, 150 (1):1–23, 1997.

Nikolet Kostur and Susana M Wadgymar. Sex allocation plasticity in response to resource and pollination availability in the annual plant brassica rapa (brassicaceae). International Journal of Plant Sciences, 184(7):499–506, 2023.

Yun-Song Lai, Di Shen, Wei Zhang, Xiaohui Zhang, Yang Qiu, Haiping Wang, Xinxin Dou, Sigeng Li, Yuanqi Wu, Jiangping Song, et al. Temperature and photoperiod changes affect cucumber sex expression by different epigenetic regulations. BMC plant biology, 18(1):268, 2018.

Fanny Laugier, Pierre Saumitou-Laprade, Philippe Vernet, Jacques Lepart, P-O Cheptou, and Mathilde Dufaÿ. Male fertility advantage within and between seasons in the perennial androdioecious plant phillyrea angustifolia. Annals of Botany, 132(7):1219–1232, 2023.

Thomas Lesaffre, John R Pannell, and Charles Mullon. An explanation for the prevalence of xy over zw sex determination in species derived from hermaphroditism. Proceedings of the National Academy of Sciences, 121(40):e2406305121, 2024.

David G Lloyd. Sexual strategies in plants iii. a quantitative method for describing the gender of plants. New Zealand journal of botany, 18(1):103–108, 1980.

Guillaume Martin and Lionel Roques. The nonstationary dynamics of fitness distributions: Asexual model with epistasis and standing variation. Genetics, 204(4):1541–1558, 2016. doi: 10.1534/genetics.116.187385.

Sandrine Maurice and Theodore H Fleming. The effect of pollen limitation on plant reproductive systems and the maintenance of sexual polymorphisms. Oikos, pages 55–60, 1995.

Sandrine Maurice, Christine Desfeux, Agnes Mignot, and Jean-Pierre Henry. Is silene acaulis (caryophyllaceae) a trioecious species? reproductive biology of two subspecies. Canadian journal of botany, 76 (3):478–485, 1998.

Susan J Mazer, Véronique A Delesalle, and Paul R Neal. Responses of floral traits to selection on primary sexual investment in spergularia marina: the battle between the sexes. Evolution, 53(3):717–731, 1999.

Sepideh Mirrahimi and Sylvain Gandon. Evolution of specialization in heterogeneous environments: equilibrium between selection, mutation and migration. Genetics, 214(2):479–491, 2020.

Laura E Perry, John R Pannell, and Marcel E Dorken. Two’s company, three’s a crowd: experimental evaluation of the evolutionary maintenance of trioecy in mercurialis annua (euphorbiaceae). PLoS One, 7(4):e35597, 2012.

Pierre Saumitou-Laprade, Philippe Vernet, Arnaud Dowkiw, Sylvain Bertrand, Sylvain Billiard, Béatrice Albert, Pierre-Henri Gouyon, and Mathilde Dufay. Polygamy or subdioecy? the impact of diallelic self-incompatibility on the sexual system in fraxinus excelsior (oleaceae). Proceedings of the Royal Society B: Biological Sciences, 285(1873):20180004, 2018.

Lukas Schärer. Tests of sex allocation theory in simultaneously hermaphroditic animals. Evolution, 63 (6):1377–1405, 2009.

RF Shaw and JD Mohler. The selective significance of the sex ratio. The American Naturalist, 87(837): 337–342, 1953.

Kohta Yoshida, Hanh Witte, Ryo Hatashima, Simo Sun, Taisei Kikuchi, Waltraud Röseler, and Ralf J Sommer. Rapid chromosome evolution and acquisition of thermosensitive stochastic sex determination in nematode androdioecious hermaphrodites. Nature Communications, 15(1):9649, 2024.

